# Genetically regulated gene expression underlies lipid traits in Hispanic cohorts

**DOI:** 10.1101/507905

**Authors:** Angela Andaleon, Lauren S. Mogil, Heather E. Wheeler

## Abstract

Plasma lipid levels are risk factors for cardiovascular disease, a leading cause of death worldwide. While many studies have been conducted in genetic variation underlying lipid levels, they mainly comprise individuals of European ancestry and thus their transferability to non-European populations is unclear. We performed genome-wide (GWAS) and imputed transcriptome-wide association studies of four lipid traits in the Hispanic Community Health Study/Study of Latinos cohort (HCHS/SoL, n = 11,103), replicated top hits in the Multi-Ethnic Study of Atherosclerosis (MESA, n = 3,855), and compared the results to the larger, predominantly European ancestry meta-analysis by the Global Lipids Genetics Consortium (GLGC, n = 196,475). In our GWAS, we found significant SNP associations in regions within or near known lipid genes, but in our admixture mapping analysis, we did not find significant associations between local ancestry and lipid phenotypes. In the imputed transcriptome-wide association study in multiple tissues and in different ethnicities, we found 59 significant gene-tissue-phenotype associations (P < 3.61×10^−8^) with 14 unique significant genes, many of which occurred across multiple phenotypes, tissues, and ethnicities and replicated in MESA (45/59) and in GLGC (44/59). These include well-studied lipid genes such as *SORT1*, *CETP*, and *PSRC1*, as well as genes that have been implicated in cardiovascular phenotypes, such as *CCL22* and *ICAM1*. The majority (40/59) of significant associations colocalized with expression quantitative trait loci (eQTLs), indicating a possible mechanism of gene regulation in lipid level variation. To fully characterize the genetic architecture of lipid traits in diverse populations, larger studies in non-European ancestry populations are needed.

## Introduction

Lipid levels are a major risk factor for cardiovascular disease, the leading cause of death in the United States [1]. While lipid levels are known to have a highly heritable component, lipid levels as a complex trait are increasingly concerning due to the growing global rate of obesity caused by rapid urbanization and high-fat foods [2]. Hispanic populations have been especially affected by this shift as Hispanic children and adolescents have the highest rate of obesity among ethnicities in the United States [1, 3]. However, like many other genetic trait studies, large lipid meta-analyses, such as the Global Lipids Genetics Consortium (GLGC), acquire information predominantly from Europeans, and these within-European discoveries may not extrapolate to other populations [4–6].

To increase our understanding of the genetic architecture of lipid traits in non-European populations, we chose to study the Hispanic Community Health Study/Study of Latinos (HCHS/SoL) [7, 8]. Phenotypes under investigation include total cholesterol (CHOL), high density lipoproteins (HDL), triglycerides (TRIG), and low density lipoproteins (LDL). This cohort has been previously studied in a GWAS for lipid traits and there were no novel loci found that replicated in independent cohorts [9].

Here, we expand upon GWAS by integrating eQTL data to predict transcriptomes in multiple tissues and in multiple ethnicities for HCHS/SoL and replication cohorts, the Multiethnic Study of Atherosclerosis (MESA) and GLGC, to further investigate the biological effects of these variants. We performed a linear mixed model GWAS for each of the four lipid phenotypes [10] in HCHS/SoL. We also performed imputed transcriptome-based association studies with PrediXcan [11] for each phenotype and cohort using gene expression prediction models built with data from 44 tissues in the Genotype-Tissue Expression Project (GTEx) [12, 13] and monocytes in MESA [14]. We calculated colocalization over all GWAS results with GTEx and MESA eQTL data, indicating possible mechanisms of action through gene regulation [13, 15]. To fully characterize the genetic architecture of traits in diverse populations, both larger transcriptome and GWAS cohorts in diverse populations are needed. All scripts used for analyses are available at https://github.com/WheelerLab/px_his_chol.

## Results

### Hispanic populations have diverse genetic ancestry between and within self-identified regions

We sought to understand the genetically regulated architecture underlying lipid traits in Hispanic populations. The genetic diversity within HCHS/SoL and other Hispanic populations has been extensively described previously and thus we concentrated on calculating cohort-specific principal components to be used as covariates in our analyses [16–18]. We calculated relatedness and genotypic principal components (PCs) with the software KING, which is designed to estimate kinship coefficients in the presence of population structure [19]. Most HCHS/SoL participants included in the analyses reported a self-identified region of ancestry in the Americas, which we included as a covariate in the regression analyses. Not only is the whole HCHS/SoL cohort genetically differentiated and heterogenous, but there is also great diversity within each self-identified region. Most individuals from the same regions tend to cluster in the same direction until the fifth principal component, but have a wide variation in eigenscores (Fig 1 and S1 Fig). In our further analyses, we used 5 PCs as fixed effects as previously performed (S2 Fig) [16].

**Figure 1.**
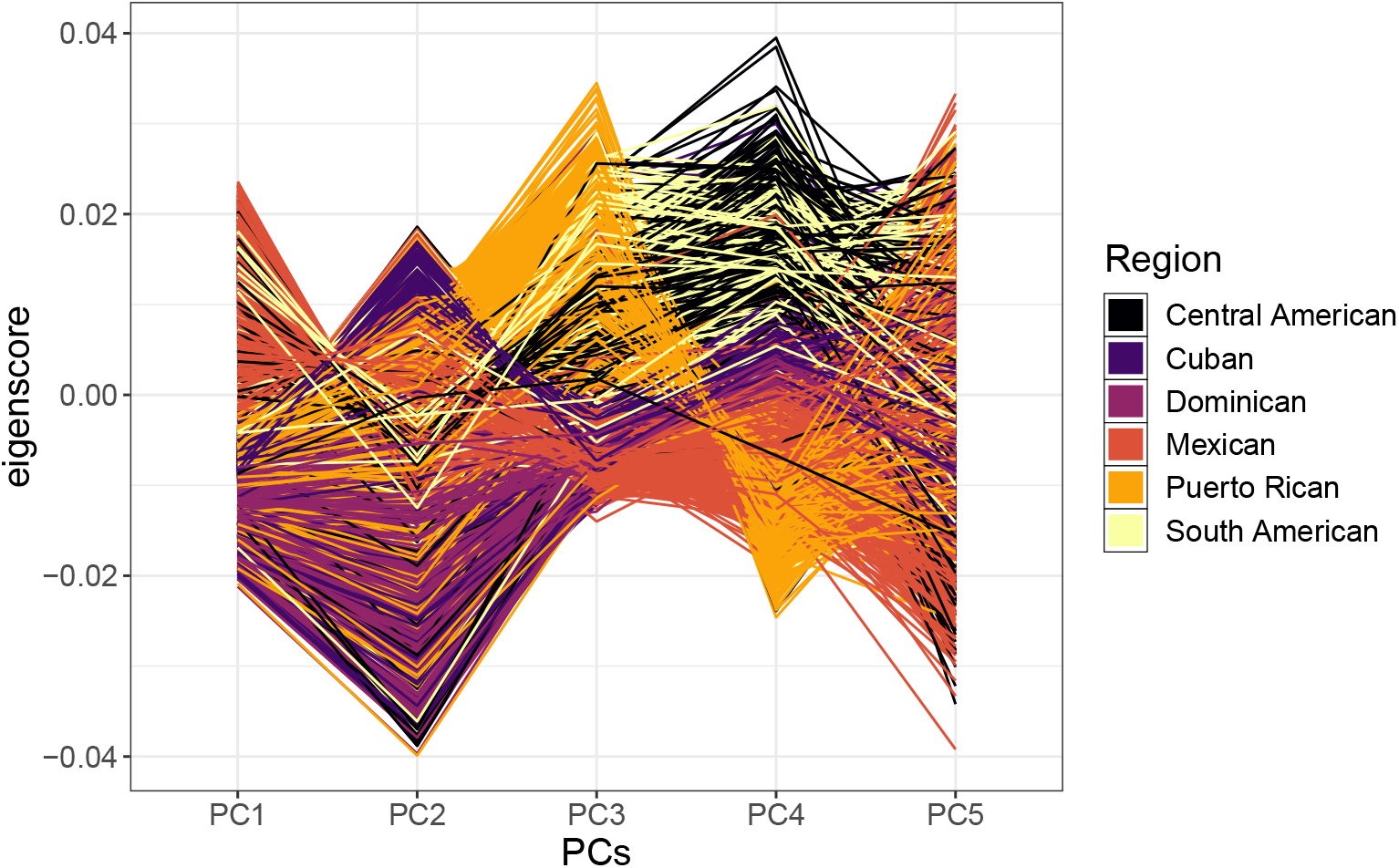
Genotypic principal component eigenscores of HCHS/SoL participants by self-identified region. Each line represents an individual in HCHS/SoL connected by their eigenscores calculated in KING and colored by self-identified region. Hispanic populations are mainly admixed between Native American, West African, and European populations, resulting in a genetically diverse and structured cohort under the umbrella term “Hispanic”.

### Joint analysis of GWAS results implicates 24 loci in lipid traits

We performed GWAS using a linear mixed model implemented in the software GEMMA [10] to investigate individual SNP associations with CHOL, HDL, TRIG, and LDL in HCHS/SoL (n = 11,103). There was little test statistic inflation among the GWAS results (S3 Fig). We followed the initial GWAS with fine-mapping by conditional and joint analyses using GCTA-COJO software [20, 21]. We used linkage disequilibrium (LD) calculated with the HCHS/SoL genotypes. We report GTCA-COJO joint effect sizes (bJ) and p-values (pJ) to emphasize independent loci signals.

In the fine-mapping joint analysis, we found multiple significant, independent SNPs associated with the four phenotypes. These SNPs include 12 with CHOL, 7 with HDL, 0 with TRIG, and 10 with LDL, totalling 29 associations (S1 Table). There were 24 unique independent SNPs across the phenotypes with some associations replicating in CHOL and an additional lipid trait (S1 Table).

Two of the significant SNPs, both associated with CHOL, had MAF < 0.01 in European populations, but both had MAF > 0.05 within HCHS/SoL (Table 1, Fig 2). rs117961479 is in LD (1000G AMR *r*^2^ = 0.632) with rs199768142, which is implicated in total cholesterol levels [22]. SNPs in LD with rs17041688 (1000G AMR *r*^2^ > 0.6) in the GWAS catalog had one association with marginal zone lymphoma, and no linked SNPs had obesity or cardiovascular associations [23].

**Table 1.** Significant conditional and joint analysis SNPs with rare allele frequency in European populations. In the GWAS of HCHS/SoL, we found 29 independent associations between SNPs and 4 lipid traits in the joint analysis, including 2 SNPs with minor allele frequency < 0.01 in Europeans. bJ indicates the effect size and pJ represents the p-value, both from a joint analysis of all the SNPs at the significant loci. The SNPs included in this table have pJ < 5×10^−8^ and EUR MAF < 0.01. Neither of these SNPs were present in our genotype data for MESA. Full results for all conditional and joint analyses are in S1 Table.

| Chr. | BP | Pheno. | SNP | bJ | pJ | GLGC P | HCHS/SoL MAF | EUR MAF | Intronic gene |
| --- | --- | --- | --- | --- | --- | --- | --- | --- | --- |
| 2 | 21199426 | CHOL | rs17041688 | -0.114 | $4.8 \times 10^{-9}$ | 0.651 | 0.112 | 0.003 | NA |
| 19 | 19410750 | CHOL | rs117961479 | -0.153 | $1.5 \times 10^{-9}$ | 0.740 | 0.060 | 0.003 | *SUGP1* |

**Figure 2.**
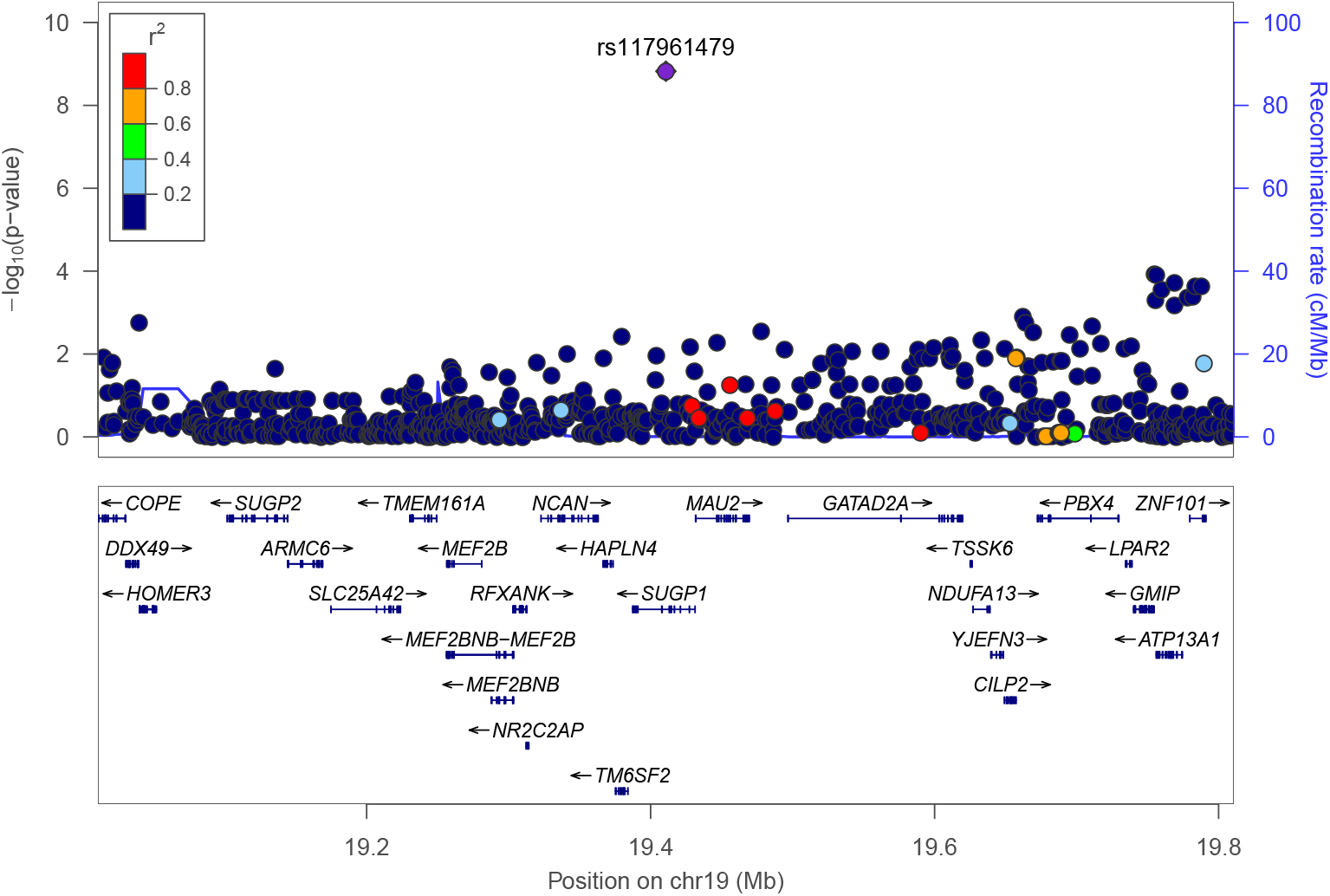
Conditional association of rs117961479 with CHOL in HCHS/SoL. Conditional and joint analysis (COJO) [21] of the HCHS/SoL CHOL GWAS results revealed a genome-wide significant signal for rs117961479 (pJ = 1.5×10^−9^). Plotted p-values for other SNPs in the region are from a conditional GWAS where the rs117961479 genotype was used as a covariate. The color of each dot represents the SNP’s linkage disequilibrium (LD) *r*^2^ with the labeled SNP in the 1000 Genomes American populations. Many nearby SNPs within or near *SUGP1* and *MAU2* are in high LD with rs117961479. These conditional results indicate one signal in this region, which was plotted with LocusZoom [24].

Percent variance explained (PVE) by all genotypes as calculated in GEMMA for each phenotype were similar to those previously observed [25, 26]. These values are CHOL: 0.269 ± 0.029, HDL: 0.180 ± 0.031, TRIG: 0.135 ± 0.030, and LDL: 0.278 ± 0.030.

### Admixture mapping does not identify ancestral associations with lipid phenotypes

Admixture mapping has been previously used in HCHS/SoL to uncover ancestral tracts, especially of Native American ancestry, that may affect traits [27, 28]. We performed admixture mapping using local ancestry estimates of European, African, and Native American chromosome tracts in each individual. We used RFMix to estimate how many alleles at each SNP came from each ancestral population [29]. We ran a separate linear mixed model for each of the three ancestries testing the number of estimated alleles from the origin ancestry for association with each phenotype in GEMMA [10]. Our reference panels for local ancestry were Iberian in Spain (IBS, n = 107), Native American (NAT, n = 27), and Yoruba in Ibadan, Nigeria (YRI, n = 108) with procedures used for reference population selection detailed in the Methods. We used a significance threshold for admixture mapping tests of P < 5.7×10^−5^, previously determined within HCHS/SoL [27]. We restricted analyses to chromosomes 16 and 19 due to their known importance in lipid traits. None of the significant SNPs within our admixture analyses reached previously determined significance. The most significant ancestry tract was on chromosome 19 from 49945850 bp to 50108983 bp for TRIG in Native American ancestry (P = 3.1×10^−4^). We include our top 1000 most significant SNPs (P < 5.3×10^−3^) in admixture analyses in S2 Table.

### Imputed transcriptome-based association study in HCHS/SoL implicates 14 genes in lipid traits

We performed imputed transcriptome-based association studies to investigate the associations of genetically predicted gene expression with the four lipid traits while also accounting for relatedness and structure in the discovery cohort HCHS/SoL (n = 11,103) [10, 11]. We used two main sets of prediction models: GTEx V6 and MESA. GTEx V6 predictor models include 44 individual tissue models built in predominantly European-American individuals (predictor population n > 70, 85% European and 15% African-American), and MESA predictor models include monocyte models from 5 populations comprising multiple ethnicities, including African-American and Hispanic (predictor population n > 233) [12–14, 30]. All GTEx results were filtered to those with green flags as described on http://predictdb.org/. We defined discovery significance as P < 3.1×10^−8^, which is 0.05/(all associations tested). Primary significance may be stringent due to the amount of eQTL sharing between transcriptomic models [31]. We tested significant associations for replication in the genotyped MESA cohort (n = 3,855) and using GWAS summary statistics from GLGC (n = 196,475) [5, 13, 32]. There were no Hispanic/Latino populations included in the GLGC analysis [5]. There was little test statistic inflation among the PrediXcan results for HCHS/SoL (S4 Fig), MESA, or GLGC. Full PrediXcan results for all cohorts are available at https://github.com/WheelerLab/px_his_chol.

Across 4 phenotypes, 44 GTEx models, and 5 MESA models, we found 59 significant gene-tissue-phenotype associations, including 14 unique significant genes. These include well-studied lipid genes such as *APOB*, *PSRC1*, *SORT1*, and *CETP* and genes that have been implicated in cardiovascular traits such as *CCL22* and *ICAM1* (Table 2, Fig 3) [33–35]. 45/59 associations replicated in MESA and 44/59 associations replicated in GLGC (P < 0.05). The only associations that did not replicate in GLGC were those that were not predicted at all due to lack of SNP overlap between the GLGC GWAS summary results and the expression prediction models. All 44 of the GLGC associations also met the more stringent threshold of P < 8.5×10^−4^, the Bonferroni adjustment for 59 tests. Despite being just 2% of the size of GLGC, 40 of the MESA associations also met the Bonferroni adjusted threshold (Table 2).

**Table 2.** Significant PrediXcan associations in HCHS/SoL and replication in MESA and GLGC. We performed PrediXcan for four lipid phenotypes in HCHS/SoL using GTEx and MESA monocyte gene expression prediction models [12–14]. In total, there are 14 unique genes across 59 significant gene-tissue-phenotype associations. MESA population abbreviations: African-American (AFA), European (CAU), Hispanic (HIS), AFA and HIS (AFHI), and all populations combined (ALL).

| Chr. | BP | Gene name | Pheno. | Model | HCHS $\beta$ | HCHS P | MESA $\beta$ | MESA P | GLGC $\beta$ | GLGC P |
| --- | --- | --- | --- | --- | --- | --- | --- | --- | --- | --- |
| 1 | 109792641 | *CELSR2* | CHOL | Esophagus Mucosa | -0.311 | $9.323 \times 10^{-12}$ | -0.424 | $3.095 \times 10^{-7}$ | -0.321 | $6.300 \times 10^{-145}$ |
| 1 | 109792641 | *CELSR2* | CHOL | Liver | -0.179 | $3.471 \times 10^{-25}$ | NA | NA | -0.123 | $2.080 \times 10^{-119}$ |
| 1 | 109792641 | *CELSR2* | CHOL | Muscle Skeletal | -0.195 | $1.526 \times 10^{-28}$ | -0.209 | $1.317 \times 10^{-9}$ | -0.144 | $2.169 \times 10^{-149}$ |
| 1 | 109792641 | *CELSR2* | CHOL | Skin Sun Exposed Lower leg | -0.216 | $1.268 \times 10^{-10}$ | -0.335 | $1.930 \times 10^{-6}$ | -0.127 | $5.802 \times 10^{-42}$ |
| 1 | 109792641 | *CELSR2* | LDL | Esophagus Mucosa | -0.388 | $2.662 \times 10^{-15}$ | -0.418 | $1.403 \times 10^{-7}$ | -0.399 | $6.393 \times 10^{-204}$ |
| 1 | 109792641 | *CELSR2* | LDL | Liver | -0.218 | $4.283 \times 10^{-32}$ | NA | NA | -0.161 | $3.712 \times 10^{-183}$ |
| 1 | 109792641 | *CELSR2* | LDL | Muscle Skeletal | -0.235 | $2.534 \times 10^{-35}$ | -0.234 | $1.500 \times 10^{-12}$ | -0.184 | $5.141 \times 10^{-220}$ |
| 1 | 109792641 | *CELSR2* | LDL | Skin Sun Exposed Lower leg | -0.274 | $3.415 \times 10^{-14}$ | -0.369 | $4.634 \times 10^{-8}$ | -0.162 | $6.032 \times 10^{-60}$ |
| 1 | 109822178 | *PSRC1* | CHOL | MESA monocytes AFA | -0.103 | $1.165 \times 10^{-11}$ | -0.210 | $1.801 \times 10^{-4}$ | -0.163 | $7.152 \times 10^{-61}$ |
| 1 | 109822178 | *PSRC1* | CHOL | MESA monocytes AFHI | -0.080 | $6.403 \times 10^{-16}$ | -0.124 | $5.406 \times 10^{-4}$ | -0.175 | $1.039 \times 10^{-102}$ |
| 1 | 109822178 | *PSRC1* | CHOL | MESA monocytes ALL | -0.082 | $1.145 \times 10^{-16}$ | -0.140 | $7.278 \times 10^{-5}$ | -0.148 | $2.074 \times 10^{-100}$ |
| 1 | 109822178 | *PSRC1* | CHOL | MESA monocytes CAU | -0.092 | $1.984 \times 10^{-16}$ | -0.188 | $5.741 \times 10^{-6}$ | -0.161 | $9.507 \times 10^{-99}$ |
| 1 | 109822178 | *PSRC1* | CHOL | MESA monocytes HIS | -0.115 | $2.185 \times 10^{-17}$ | -0.247 | $1.842 \times 10^{-6}$ | -0.249 | $5.981 \times 10^{-133}$ |
| 1 | 109822178 | *PSRC1* | CHOL | Esophagus Mucosa | -0.173 | $2.883 \times 10^{-15}$ | -0.226 | $3.018 \times 10^{-8}$ | -0.149 | $1.324 \times 10^{-131}$ |
| 1 | 109822178 | *PSRC1* | CHOL | Liver | -0.142 | $3.098 \times 10^{-23}$ | NA | NA | -0.105 | $7.243 \times 10^{-130}$ |
| 1 | 109822178 | *PSRC1* | CHOL | Lung | -0.344 | $3.267 \times 10^{-8}$ | -0.316 | $1.417 \times 10^{-2}$ | NA | NA |
| 1 | 109822178 | *PSRC1* | CHOL | Muscle Skeletal | -0.267 | $7.747 \times 10^{-9}$ | -0.438 | $4.135 \times 10^{-6}$ | -0.235 | $5.258 \times 10^{-67}$ |
| 1 | 109822178 | *PSRC1* | CHOL | Pancreas | -0.204 | $3.887 \times 10^{-10}$ | -0.457 | $3.811 \times 10^{-9}$ | -0.161 | $8.042 \times 10^{-84}$ |
| 1 | 109822178 | *PSRC1* | CHOL | Pituitary | -0.196 | $1.913 \times 10^{-11}$ | -0.421 | $1.125 \times 10^{-9}$ | -0.149 | $3.348 \times 10^{-85}$ |
| 1 | 109822178 | *PSRC1* | CHOL | Testis | -0.211 | $2.429 \times 10^{-15}$ | -0.235 | $2.543 \times 10^{-6}$ | -0.162 | $2.286 \times 10^{-94}$ |
| 1 | 109822178 | *PSRC1* | CHOL | Whole Blood | -0.822 | $1.722 \times 10^{-24}$ | -0.901 | $3.215 \times 10^{-9}$ | -0.691 | $3.309 \times 10^{-175}$ |
| 1 | 109822178 | *PSRC1* | LDL | MESA monocytes AFA | -0.127 | $6.119 \times 10^{-15}$ | -0.196 | $2.732 \times 10^{-4}$ | -0.210 | $3.754 \times 10^{-90}$ |
| 1 | 109822178 | *PSRC1* | LDL | MESA monocytes AFHI | -0.095 | $2.221 \times 10^{-19}$ | -0.125 | $2.602 \times 10^{-4}$ | -0.223 | $1.057 \times 10^{-151}$ |
| 1 | 109822178 | *PSRC1* | LDL | MESA monocytes ALL | -0.099 | $1.499 \times 10^{-20}$ | -0.139 | $4.357 \times 10^{-5}$ | -0.187 | $4.240 \times 10^{-147}$ |
| 1 | 109822178 | *PSRC1* | LDL | MESA monocytes CAU | -0.112 | $1.179 \times 10^{-20}$ | -0.184 | $3.897 \times 10^{-6}$ | -0.206 | $2.933 \times 10^{-146}$ |
| 1 | 109822178 | *PSRC1* | LDL | MESA monocytes HIS | -0.138 | $3.418 \times 10^{-21}$ | -0.245 | $8.567 \times 10^{-7}$ | -0.315 | $7.900 \times 10^{-194}$ |
| 1 | 109822178 | *PSRC1* | LDL | Esophagus Mucosa | -0.214 | $7.540 \times 10^{-20}$ | -0.244 | $4.273 \times 10^{-10}$ | -0.189 | $1.019 \times 10^{-193}$ |
| 1 | 109822178 | *PSRC1* | LDL | Liver | -0.173 | $2.231 \times 10^{-29}$ | NA | NA | -0.135 | $1.613 \times 10^{-190}$ |
| 1 | 109822178 | *PSRC1* | LDL | Lung | -0.451 | $1.458 \times 10^{-11}$ | -0.364 | $3.239 \times 10^{-3}$ | NA | NA |
| 1 | 109822178 | *PSRC1* | LDL | Muscle Skeletal | -0.341 | $6.847 \times 10^{-12}$ | -0.402 | $1.040 \times 10^{-5}$ | -0.277 | $1.836 \times 10^{-87}$ |
| 1 | 109822178 | *PSRC1* | LDL | Pancreas | -0.262 | $6.506 \times 10^{-14}$ | -0.480 | $1.079 \times 10^{-10}$ | -0.201 | $2.633 \times 10^{-121}$ |
| 1 | 109822178 | *PSRC1* | LDL | Pituitary | -0.254 | $5.485 \times 10^{-16}$ | -0.442 | $2.330 \times 10^{-11}$ | -0.183 | $1.146 \times 10^{-120}$ |
| 1 | 109822178 | *PSRC1* | LDL | Skin Not Sun Exposed Suprapubic | -0.187 | $1.492 \times 10^{-10}$ | -0.228 | $2.529 \times 10^{-4}$ | NA | NA |
| 1 | 109822178 | *PSRC1* | LDL | Testis | -0.255 | $5.398 \times 10^{-19}$ | -0.236 | $8.441 \times 10^{-7}$ | -0.208 | $4.254 \times 10^{-139}$ |
| 1 | 109822178 | *PSRC1* | LDL | Whole Blood | -1.011 | $1.096 \times 10^{-31}$ | -0.959 | $4.887 \times 10^{-11}$ | -0.866 | $7.059 \times 10^{-253}$ |
| 1 | 109852192 | *SORT1* | CHOL | Liver | -0.166 | $5.472 \times 10^{-28}$ | NA | NA | -0.106 | $1.046 \times 10^{-123}$ |
| 1 | 109852192 | *SORT1* | LDL | Liver | -0.203 | $1.217 \times 10^{-35}$ | NA | NA | -0.136 | $7.449 \times 10^{-183}$ |
| 1 | 109852192 | *SORT1* | LDL | Pancreas | -0.284 | $3.414 \times 10^{-9}$ | -0.300 | $1.318 \times 10^{-4}$ | -0.237 | $6.478 \times 10^{-96}$ |
| 1 | 110026101 | *ATXN7L2* | LDL | Liver | -0.103 | $2.109 \times 10^{-9}$ | NA | NA | -0.064 | $6.624 \times 10^{-53}$ |
| 1 | 110230436 | *GSTM1* | CHOL | Small Intestine Terminal Ileum | -4.469 | $1.481 \times 10^{-8}$ | NA | NA | NA | NA |
| 1 | 110230436 | *GSTM1* | LDL | Small Intestine Terminal Ileum | -5.404 | $1.747 \times 10^{-10}$ | NA | NA | NA | NA |
| 2 | 21224301 | *APOB* | CHOL | Adipose Subcutaneous | 0.193 | $2.400 \times 10^{-11}$ | 0.199 | $6.413 \times 10^{-4}$ | 0.166 | $2.972 \times 10^{-67}$ |
| 2 | 21224301 | *APOB* | LDL | Adipose Subcutaneous | 0.233 | $6.653 \times 10^{-14}$ | 0.209 | $1.854 \times 10^{-4}$ | 0.193 | $1.117 \times 10^{-86}$ |
| 15 | 58702768 | *LIPC* | HDL | Liver | -0.175 | $2.558 \times 10^{-9}$ | NA | NA | NA | NA |
| 16 | 56995762 | *CETP* | HDL | MESA monocytes AFHI | -0.425 | $7.337 \times 10^{-30}$ | -0.560 | $8.862 \times 10^{-8}$ | -0.951 | 0 |
| 16 | 56995762 | *CETP* | HDL | MESA monocytes ALL | -0.259 | $1.076 \times 10^{-27}$ | -0.327 | $2.156 \times 10^{-7}$ | -0.518 | 0 |
| 16 | 56995762 | *CETP* | HDL | MESA monocytes CAU | -0.202 | $7.302 \times 10^{-}$ | -0.408 | $6.857 \times 10^{-7}$ | -0.675 | 0 |
| 16 | 56995762 | *CETP* | HDL | MESA monocytes HIS | -0.290 | $1.848 \times 10^{-31}$ | -0.572 | $8.532 \times 10^{-11}$ | -0.835 | 0 |
| 16 | 56995762 | *CETP* | HDL | Artery Coronary | -0.454 | $1.286 \times 10^{-27}$ | -0.983 | $1.372 \times 10^{-13}$ | NA | NA |
| 16 | 56995762 | *CETP* | HDL | Cells Transformed fibroblasts | -0.120 | $1.504 \times 10^{-8}$ | -0.144 | $1.093 \times 10^{-1}$ | NA | NA |
| 16 | 56995762 | *CETP* | HDL | Esophagus Mucosa | -1.011 | $6.544 \times 10^{-32}$ | -2.084 | $2.853 \times 10^{-14}$ | NA | NA |
| 16 | 56995762 | *CETP* | HDL | Lung | -0.477 | $1.515 \times 10^{-24}$ | -0.979 | $2.270 \times 10^{-12}$ | NA | NA |
| 16 | 57023397 | *NLRC5* | HDL | Cells Transformed fibroblasts | -0.107 | $3.175 \times 10^{-18}$ | -0.218 | $8.743 \times 10^{-4}$ | NA | NA |
| 16 | 57023397 | *NLRC5* | HDL | Heart Atrial Appendage | 0.156 | $1.643 \times 10^{-9}$ | 0.062 | $5.211 \times 10^{-1}$ | NA | NA |
| 16 | 57392684 | *CCL22* | HDL | Esophagus Muscularis | -9.841 | $1.405 \times 10^{-38}$ | -15.845 | $3.502 \times 10^{-14}$ | NA | NA |
| 19 | 10381511 | *ICAM1* | HDL | Esophagus Muscularis | -0.181 | $1.998 \times 10^{-8}$ | -0.439 | $1.021 \times 10^{-2}$ | -0.153 | $1.068 \times 10^{-10}$ |
| 19 | 11039409 | *TIMM29* | CHOL | Brain Cerebellar Hemisphere | 5.137 | $1.754 \times 10^{-13}$ | NA | NA | NA | NA |
| 19 | 11309971 | *DOCK6* | HDL | Artery Tibial | 0.457 | $2.930 \times 10^{-9}$ | 0.776 | $1.015 \times 10^{-3}$ | 0.268 | $2.196 \times 10^{-5}$ |
| 19 | 44645710 | *ZNF234* | LDL | Breast Mammary Tissue | -0.171 | $1.872 \times 10^{-8}$ | 0.265 | $1.669 \times 10^{-3}$ | NA | NA |

**Figure 3.**
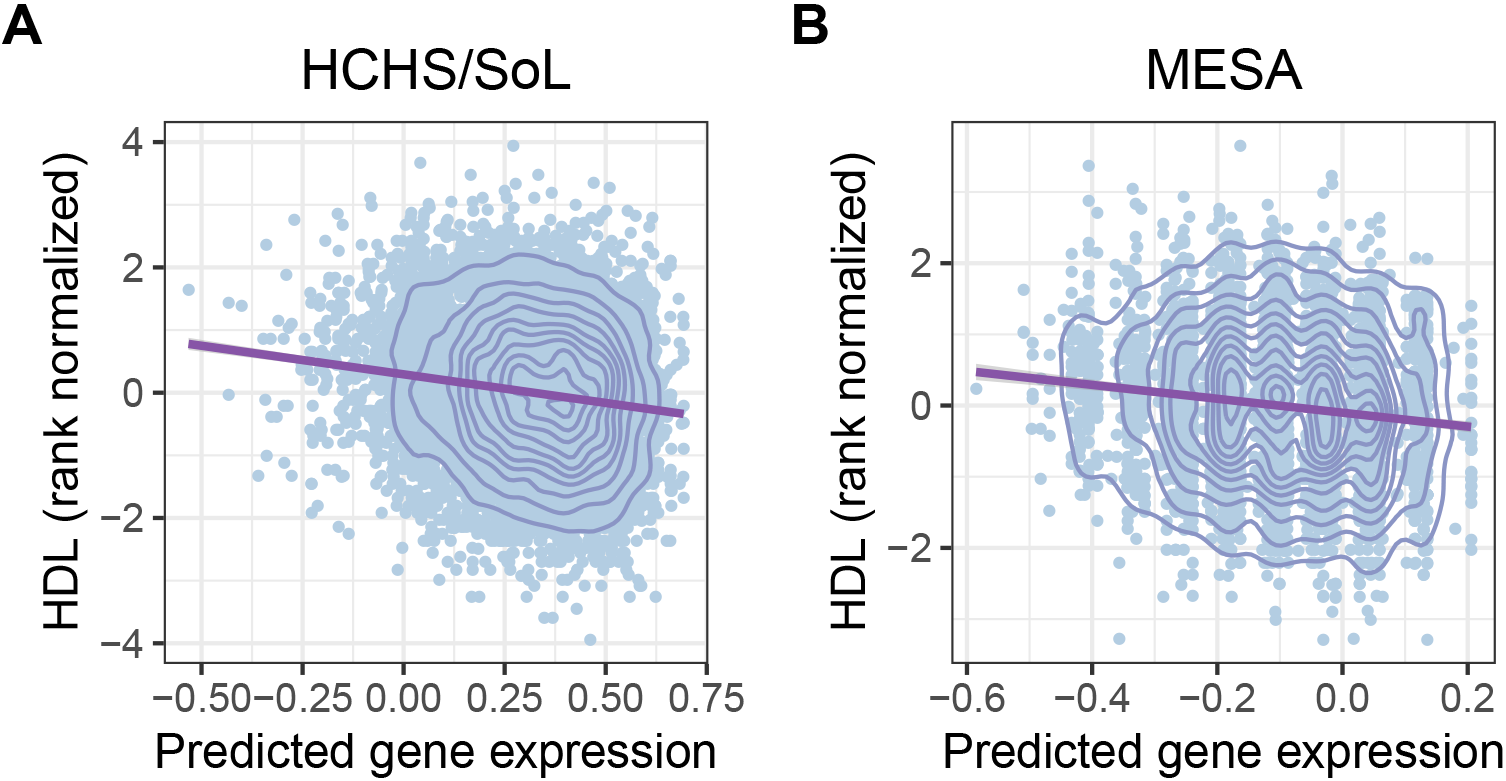
Predicted expression vs. observed phenotype for *CETP*. Using the GTEx Artery Coronary expression prediction model, increased predicted gene expression of *CETP* is significantly associated with decreased observed HDL in both HCHS/SoL (A) and MESA (B) (Table 2), which is consistent with previous studies of *CETP* [35].

Though most of the results were from the 44 predominantly European GTEx tissue models, we also had significant PrediXcan results using the five MESA monocyte models built in various ethnicities [14]. 14/59 of the significant associations in HCHS/SoL were in MESA models, with 4 associations in *CETP* and 10 associations in *PSRC1* in the same effect direction as the other tissues (Table 2).

A recent study investigated cardiometabolic traits, including the four lipid phenotypes, within diverse cohorts using PrediXcan GTEx V7 models, which, unlike V6 used in our analyses, include only European ancestry individuals [36]. They found 12 gene-lipid-tissue associations at P < 2.6×10^−5^, one of which replicated in GLGC. We compared their novel GTEx tissue results to our results. None of their gene associations replicated in our analyses (all P > 0.09).

### A majority of significant PrediXcan gene associations have colocalized GWAS and eQTL signals

We further investigated whether the PrediXcan gene associations had colocalized signal with known eQTLs. Colocalization provides additional evidence that the SNPs in a given expression model are functioning via gene regulation to affect lipid traits. We restricted variants to those within PrediXcan models and we tested them for colocalization of eQTLs and lipid trait GWAS associations. We used the software COLOC, which estimates the colocalization probability for a SNP between an eQTL and a GWAS hit [15]. We have previously applied COLOC to gene-level PrediXcan results and observed clustering of significant genes into non-colocalized, colocalized, or unknown signal [13]. P3 estimations indicate the probability of independent signals from an eQTL association and a GWAS association, with P3 > 0.5 indicating no colocalization. P4 > 0.5 indicates a shared eQTL and GWAS association of variants within the prediction model, and if neither P3 nor P4 > 0.5, whether or not the GWAS and the eQTL signals are colocalized is unknown [13, 15]. We used eQTL data from GTEx V6 and MESA monocytes, the same populations as the PrediXcan predictors [14, 30].

We calculated colocalization probabilities for all significant gene-phenotype-tissue associations. Of these associations, 40/59 had a better than chance probability of colocalization between GWAS and eQTL signals (P4 > 0.5) (Table 3). This included the genes *PSRC1*, *APOB*, *CELSR2*, *SORT1*, *CETP*, *DOCK6*, *ICAM1*, *CCL22*, and *LIPC*. Of the remaining 19 associations, 12 were likely independent signals (P3 > 0.5) and 7 were inconclusive. Full results are available at https://github.com/WheelerLab/px_his_chol.

**Table 3.** Gene associations and colocalization with eQTLs. We performed COLOC to test colocalization between HCHS/SoL GWAS and eQTL data in our transcriptomic models (44 GTEx, 5 MESA). We found 40/59 of significant (P < 3.61×10^−8^) PrediXcan associations had significant probability (P4 > 0.5) for colocalization, indicating that the variants may be acting through gene regulation to affect the phenotypes. P3 > 0.5 indicates independence between GWAS and eQTL signals. Full results are available at https://github.com/WheelerLab/px_his_chol.

| Chr. | BP | Gene | Model | Pheno. | HCHS $\beta$ | HCHS P | P3 | P4 |
| --- | --- | --- | --- | --- | --- | --- | --- | --- |
| 1 | 109792641 | *CELSR2* | Liver | CHOL | -0.179 | $3.471 \times 10^{-25}$ | 0.001 | 0.999 |
| 1 | 109792641 | *CELSR2* | Muscle Skeletal | CHOL | -0.195 | $1.526 \times 10^{-28}$ | 0.001 | 0.999 |
| 1 | 109792641 | *CELSR2* | Liver | LDL | -0.218 | $4.283 \times 10^{-32}$ | 0.001 | 0.999 |
| 1 | 109792641 | *CELSR2* | Muscle Skeletal | LDL | -0.235 | $2.534 \times 10^{-35}$ | 0.002 | 0.998 |
| 1 | 109792641 | *CELSR2* | Skin Sun Exposed Lower Leg | LDL | -0.274 | $3.415 \times 10^{-14}$ | 0.006 | 0.982 |
| 1 | 109792641 | *CELSR2* | Skin Sun Exposed Lower Leg | CHOL | -0.216 | $1.268 \times 10^{-10}$ | 0.007 | 0.982 |
| 1 | 109792641 | *CELSR2* | Esophagus Mucosa | LDL | -0.388 | $2.662 \times 10^{-15}$ | 0.028 | 0.959 |
| 1 | 109792641 | *CELSR2* | Esophagus Mucosa | CHOL | -0.311 | $9.323 \times 10^{-12}$ | 0.028 | 0.959 |
| 1 | 109822178 | *PSRC1* | Liver | CHOL | -0.142 | $3.098 \times 10^{-23}$ | 0.001 | 0.999 |
| 1 | 109822178 | *PSRC1* | Liver | LDL | -0.173 | $2.231 \times 10^{-29}$ | 0.001 | 0.999 |
| 1 | 109822178 | *PSRC1* | Esophagus Mucosa | LDL | -0.214 | $7.540 \times 10^{-20}$ | 0.002 | 0.998 |
| 1 | 109822178 | *PSRC1* | Esophagus Mucosa | CHOL | -0.173 | $2.883 \times 10^{-15}$ | 0.003 | 0.997 |
| 1 | 109822178 | *PSRC1* | Pancreas | CHOL | -0.204 | $3.887 \times 10^{-10}$ | 0.010 | 0.981 |
| 1 | 109822178 | *PSRC1* | Pancreas | LDL | -0.262 | $6.506 \times 10^{-14}$ | 0.010 | 0.981 |
| 1 | 109822178 | *PSRC1* | Whole Blood | CHOL | -0.822 | $1.722 \times 10^{-24}$ | 0.019 | 0.977 |
| 1 | 109822178 | *PSRC1* | Whole Blood | LDL | -1.011 | $1.096 \times 10^{-31}$ | 0.019 | 0.977 |
| 1 | 109822178 | *PSRC1* | Skin Not Sun Exposed Suprapubic | LDL | -0.187 | $1.492 \times 10^{-10}$ | 0.044 | 0.906 |
| 1 | 109822178 | *PSRC1* | Pituitary | LDL | -0.254 | $5.485 \times 10^{-16}$ | 0.019 | 0.905 |
| 1 | 109822178 | *PSRC1* | Pituitary | CHOL | -0.196 | $1.913 \times 10^{-11}$ | 0.020 | 0.902 |
| 1 | 109822178 | *PSRC1* | Testis | CHOL | -0.211 | $2.429 \times 10^{-15}$ | 0.090 | 0.891 |
| 1 | 109822178 | *PSRC1* | Testis | LDL | -0.255 | $5.398 \times 10^{-19}$ | 0.090 | 0.891 |
| 1 | 109822178 | *PSRC1* | Muscle Skeletal | CHOL | -0.267 | $7.747 \times 10^{-9}$ | 0.070 | 0.801 |
| 1 | 109822178 | *PSRC1* | Muscle Skeletal | LDL | -0.341 | $6.847 \times 10^{-12}$ | 0.072 | 0.795 |
| 1 | 109822178 | *PSRC1* | MESA monocytes HIS | LDL | -0.138 | $3.418 \times 10^{-21}$ | 0.285 | 0.715 |
| 1 | 109822178 | *PSRC1* | MESA monocytes HIS | CHOL | -0.115 | $2.185 \times 10^{-17}$ | 0.293 | 0.707 |
| 1 | 109822178 | *PSRC1* | Lung | CHOL | -0.344 | $3.267 \times 10^{-8}$ | 0.067 | 0.437 |
| 1 | 109822178 | *PSRC1* | Lung | LDL | -0.451 | $1.458 \times 10^{-11}$ | 0.067 | 0.432 |
| 1 | 109822178 | *PSRC1* | MESA monocytes CAU | CHOL | -0.092 | $1.984 \times 10^{-16}$ | 0.998 | 0.002 |
| 1 | 109822178 | *PSRC1* | MESA monocytes CAU | LDL | -0.112 | $1.179 \times 10^{-20}$ | 0.998 | 0.002 |
| 1 | 109822178 | *PSRC1* | MESA monocytes AFHI | CHOL | -0.080 | $6.403 \times 10^{-16}$ | 0.999 | 0.001 |
| 1 | 109822178 | *PSRC1* | MESA monocytes AFHI | LDL | -0.095 | $2.221 \times 10^{-19}$ | 0.999 | 0.001 |
| 1 | 109822178 | *PSRC1* | MESA monocytes AFA | LDL | -0.127 | $6.119 \times 10^{-15}$ | 1.000 | 0.000 |
| 1 | 109822178 | *PSRC1* | MESA monocytes AFA | CHOL | -0.103 | $1.165 \times 10^{-11}$ | 1.000 | 0.000 |
| 1 | 109822178 | *PSRC1* | MESA monocytes ALL | LDL | -0.099 | $1.499 \times 10^{-20}$ | 1.000 | 0.000 |
| 1 | 109822178 | *PSRC1* | MESA monocytes ALL | CHOL | -0.082 | $1.145 \times 10^{-16}$ | 1.000 | 0.000 |
| 1 | 109852192 | *SORT1* | Liver | CHOL | -0.166 | $5.472 \times 10^{-28}$ | 0.001 | 0.999 |
| 1 | 109852192 | *SORT1* | Liver | LDL | -0.203 | $1.217 \times 10^{-35}$ | 0.001 | 0.999 |
| 1 | 109852192 | *SORT1* | Pancreas | LDL | -0.284 | $3.414 \times 10^{-9}$ | 0.035 | 0.910 |
| 1 | 110026101 | *ATXN7L2* | Liver | LDL | -0.103 | $2.109 \times 10^{-9}$ | 0.988 | 0.010 |
| 1 | 110230436 | *GSTM1* | Small Intestine Terminal Ileum | CHOL | -4.469 | $1.481 \times 10^{-8}$ | 0.080 | 0.340 |
| 1 | 110230436 | *GSTM1* | Small Intestine Terminal Ileum | LDL | -5.404 | $1.747 \times 10^{-10}$ | 0.080 | 0.340 |
| 2 | 21224301 | *APOB* | Adipose Subcutaneous | CHOL | 0.193 | $2.400 \times 10^{-11}$ | 0.167 | 0.833 |
| 2 | 21224301 | *APOB* | Adipose Subcutaneous | LDL | 0.233 | $6.653 \times 10^{-14}$ | 0.305 | 0.695 |
| 15 | 58702768 | *LIPC* | Liver | HDL | -0.175 | $2.558 \times 10^{-9}$ | 0.061 | 0.652 |
| 16 | 56995762 | *CETP* | MESA monocytes ALL | HDL | -0.259 | $1.076 \times 10^{-27}$ | 0.002 | 0.998 |
| 16 | 56995762 | *CETP* | MESA monocytes AFHI | HDL | -0.425 | $7.337 \times 10^{-30}$ | 0.002 | 0.998 |
| 16 | 56995762 | *CETP* | MESA monocytes HIS | HDL | -0.290 | $1.848 \times 10^{-31}$ | 0.005 | 0.995 |
| 16 | 56995762 | *CETP* | Esophagus Mucosa | HDL | -1.011 | $6.544 \times 10^{-32}$ | 0.007 | 0.963 |
| 16 | 56995762 | *CETP* | Lung | HDL | -0.477 | $1.515 \times 10^{-24}$ | 0.007 | 0.961 |
| 16 | 56995762 | *CETP* | Artery Coronary | HDL | -0.454 | $1.286 \times 10^{-27}$ | 0.026 | 0.819 |
| 16 | 56995762 | *CETP* | Cells Transformed Fibroblasts | HDL | -0.120 | $1.504 \times 10^{-8}$ | 0.509 | 0.124 |
| 16 | 56995762 | *CETP* | MESA monocytes CAU | HDL | -0.202 | $7.302 \times 10^{-23}$ | 0.997 | 0.003 |
| 16 | 57023397 | *NLRC5* | Heart Atrial Appendage | HDL | 0.156 | $1.643 \times 10^{-9}$ | 0.130 | 0.060 |
| 16 | 57023397 | *NLRC5* | Cells Transformed Fibroblasts | HDL | -0.107 | $3.175 \times 10^{-18}$ | 1.000 | 0.000 |
| 16 | 57392684 | *CCL22* | Esophagus Muscularis | HDL | -9.841 | $1.405 \times 10^{-38}$ | 0.034 | 0.749 |
| 19 | 10381511 | *ICAM1* | Esophagus Muscularis | HDL | -0.181 | $1.998 \times 10^{-8}$ | 0.023 | 0.802 |
| 19 | 11039409 | *TIMM29* | Brain Cerebellar Hemisphere | CHOL | 5.137 | $1.754 \times 10^{-13}$ | 0.071 | 0.086 |
| 19 | 11309971 | *DOCK6* | Artery Tibial | HDL | 0.457 | $2.930 \times 10^{-9}$ | 0.020 | 0.820 |
| 19 | 44645710 | *ZNF234* | Breast Mammary Tissue | LDL | -0.171 | $1.872 \times 10^{-8}$ | 0.095 | 0.250 |

Of the 8 significant associations found in liver, 7/8 had a significant P4 (P4 > 0.5) and 6/8 with P4 > 0.99, highlighting the liver’s importance in lipid metabolism and possibly indicating its action through gene regulation. While somewhat expected, these results are notable because the GTEx liver tissue has a smaller sample size (n = 97) than other tissues with multiple associations (whole blood: 2/49, n = 338; MESA monocytes ALL 3/49, n = 1,163), signifying that liver is an especially important tissue in lipid metabolism as extensively studied previously [15, 37].

We found differences in colocalization probabilities between the MESA models of different ethnicities. This is most prevalent in the *CETP* model, a known driver of cholesterol metabolism, where only the models including Hispanic genotypes had P4 > 0.99, while the CAU model displayed independent GWAS and eQTL signals (Table 3). This is likely due to the shared LD patterns in the Hispanic eQTL and GWAS cohorts (Fig 4). SNPs included in the MESA HIS and MESA CAU PrediXcan models for *CETP* are different because elastic net variable selection, which was used to generate the model in each population [14], is partially dependent on correlation among variables, i.e. LD. Thus, since the LD pattern of HCHS more closely resembles HIS than CAU and because linked SNPs in HCHS and HIS were most significantly associated with HDL and *CETP* expression, respectively, the colocalization probability is much higher in the HIS model than the CAU model (Fig 4, Table 3).

**Figure 4.**
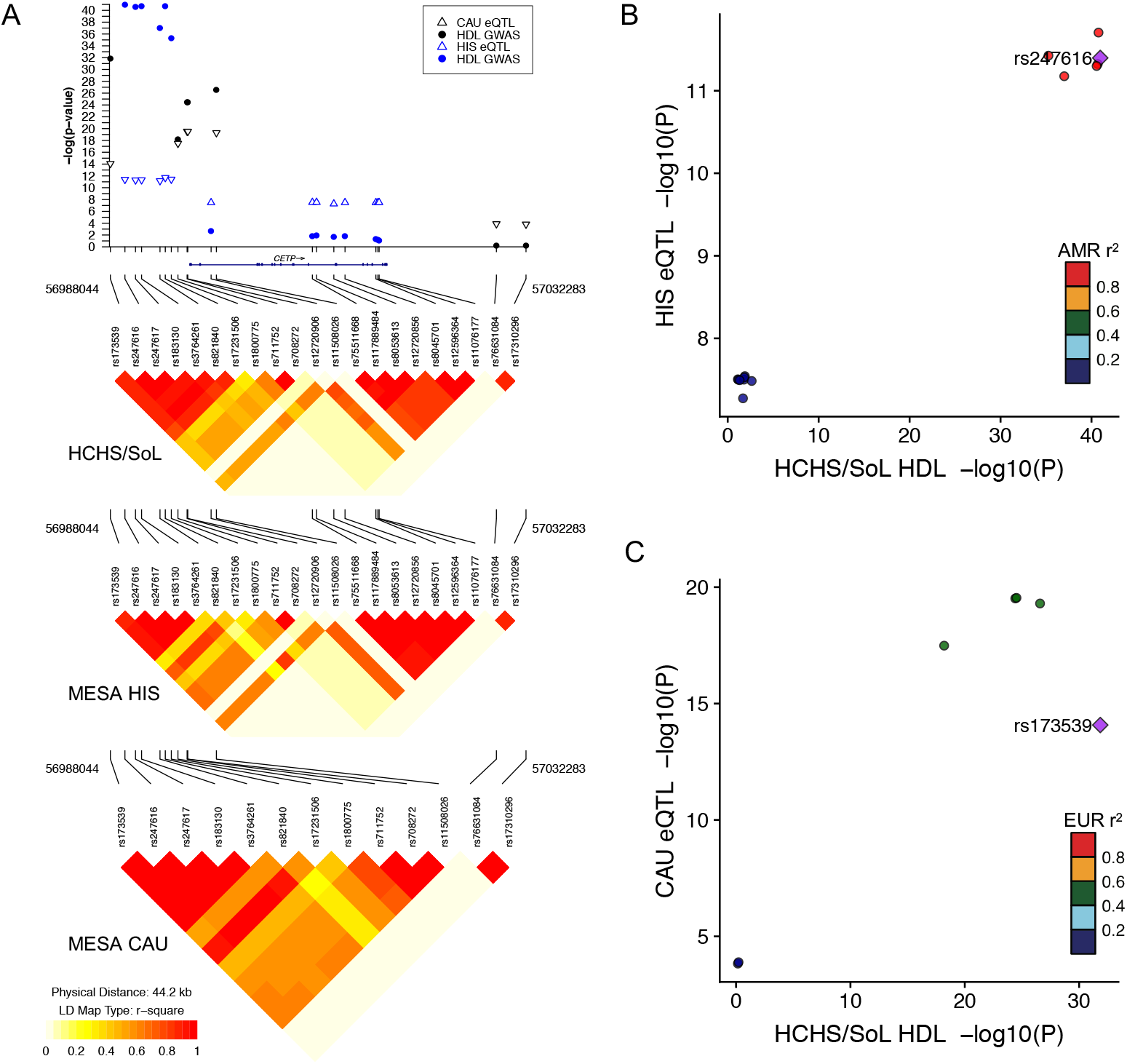
Colocalization of HCHS/SoL HDL GWAS and MESA eQTL signals at the *CETP* locus. (A) Zoomed in Manhattan plot of the key SNPs driving the *CETP* association in the MESA HIS and MESA CAU PrediXcan models. Filled circles represent HDL associations in HCHS/SoL and open triangles represent eQTLs in MESA (up- or down-triangles indicate the effect allele is associated with increased or decreased *CETP* expression, respectively). Blue symbols represent SNPs in the MESA HIS PrediXcan model and black symbols represent SNPs in the MESA CAU PrediXcan model. Linkage disequilibrium (LD) plots are labeled with the population genotypes used to calculate r^2^. Note several SNPs present in the Hispanic populations were monomorphic in MESA CAU and thus not included on the plot. Comparison between eQTL and HCHS/SoL HDL GWAS p-values for SNPs in either the MESA HIS (B) or MESA CAU (C) PrediXcan model. The most significant HCHS/SoL HDL GWAS SNP is the index SNP (purple diamond) in each plot. LD is calculated from the 1000 Genomes American (AMR) or European (EUR) populations. Note the index SNP is linked to the strongest eQTL signals in the MESA HIS model, but not in the MESA CAU model. Plots were generated with snp.plotter (A) and LocusCompare (B,C) [38, 39].

## Discussion

We integrated expression quantitative trait loci (eQTL) data from multiple tissues and multiple ethnicities in transcriptome-wide association and colocalization studies to investigate the biological mechanisms underlying lipid trait variation. GWAS have previously been performed in the Hispanic discovery cohort, HCHS/SoL, with no novel, replicable associations found [9], similar to our SNP-level findings. We acknowledge our GWAS findings were limited due to our restriction to SNPs present in PrediXcan models, which were derived from predominantly European cohorts. However, with PrediXcan, we found 59 transcriptome-wide significant gene associations with lipid traits in HCHS/SoL, with 45/59 gene-tissue-phenotype combinations replicating in MESA and 44/59 combinations replicating in GLGC. MESA and GLGC are of diverse (African-American, European, and Hispanic ancestry) and European-only ancestry, respectively, with a similar replication rate in MESA, which has < 2% of the cohort size of GLGC.

Imputed-transcriptome based association methods like PrediXcan offer benefits over SNP-level GWAS in producing actionable results [11]. Running PrediXcan across GTEx models can implicate particular tissues in a phenotype. Here, we find the most (8/59) significant associations with the GTEx liver models compared to other tissue models, which is encouraging as the liver is key in cholesterol synthesis and metabolism (Table 2, Fig 3) [37]. PrediXcan results include a direction of effect, and many of our results had the same direction of effect as *in vivo* observations in humans and mice [35, 40]. For example, we found increased expression of *PSRC1* is significantly associated with lower CHOL and LDL in multiple tissues and ethnicities. The same relationship has been previously observed in the cholesterol metabolism of mice and in the measured gene expression in an Indian population [41, 42]. Our *CETP* results indicated that higher *CETP* expression is associated with lower HDL levels, which has been previously observed extensively within humans and has become a potential drug target for preventing atherosclerosis [35, 43–46]. Directions of effect revealed by PrediXcan can help elucidate genetic pathways of metabolism or potential drug targets [11].

Our significance threshold was stringent because we used the Bonferroni correction over all associations, but many of the eQTLs between genes and tissues, especially related tissues, are the same or in linkage disequilibrium [31]. *PSRC1* and *SORT1* are linked and were both highly significant in our analyses. Knowing liver is the key tissue involved in lipid metabolism may help us prioritize the slightly more significant *SORT1* liver association over *PSRC1*, even though *PSRC1* associations were more significant in 9 other tissues (Table 3). However, because liver expression between the genes is highly correlated (Pearson R=0.75) [47], conditional analyses cannot help us distinguish between one or both genes contributing to lipid metabolism. In one functional study, overexpression and knockdown of *Sort1* in mice altered cholesterol levels, while perturbation of *Psrc1* did not [48], suggesting *SORT1* is the more likely causal gene [47]. However, subsequent functional studies also implicate PSRC1 in cholesterol transport in apoE^−/−^ mice [41]. Thus, additional evidence is often necessary to identify the most likely causal gene or genes, due to potential confounding PrediXcan results from linkage and co-regulation of gene expression [47].

We found numerous significant gene associations with lipid traits that displayed colocalization with eQTL signals and replication with GLGC (Tables 2 and 3). These include *CELSR2*, *PSRC1*, *SORT1*, *GSTM1*, *APOB*, *LIPC*, *CETP*, *NLRC5*, *CCL22*, *ICAM1*, *TIMM29*, *DOCK6*, *ZNF234*. Of these genes, *CELSR2*, *PSRC1*, *GSTM1*, *SORT1*, *APOB*, *LIPC*, *CETP*, and *DOCK6*, which all have been extensively implicated and studied in the processes of lipid metabolism in humans [5, 49–52]. Other genes within this set are located near these well-studied genes, such as *NLRC5*, which is 5,645 bp downstream from *CETP*. Here, we describe the current implications of genes that are greater than 100 kb away from the closest significant gene or previously observed known lipid gene (Table 3).

Increased *CCL22* predicted expression significantly associated with lower HDL levels, showed evidence of colocalization, and the gene is located 375 kb downstream from *CETP* (Tables 2 and 3). It is involved in immunoregulatory and inflammatory processes, and variants within it have been implicated in multiple sclerosis and lupus [53, 54]. *Apoe* knockout mice with a high cholesterol diet were found to have significantly higher CCL22 serum levels than similar mice on a regular diet, contributing to the progression of atherosclerosis [55]. A separate study in humans from the same group found increased CCL22 abundance associated with ischemic heart disease progression [56]. These observations correlate with our observed association of higher predicted *CCL22* expression with lower HDL levels, as higher amounts of HDL might protect against atherosclerosis [57].

Increased *ICAM1* predicted expression is significantly associated with lower HDL levels, it showed evidence of colocalization, and it is located 803 kb upstream of *LDLR*, a prominent lipid gene. It produces ICAM-1, an intercellular adhesion molecule. In male rats, a high cholesterol diet was found to significantly influence *ICAM-1* molecule levels in the aorta, signifying a correlation between a cholesterol diet and *ICAM-1* expression, and *ICAM-1* molecule levels were negatively correlated with baseline HDL levels [58]. Another study found that HDL suppresses expression of *ICAM1* through transfer of *microRNA-223* to endothelial cells, which correlated with our observed association [59].

Increased *GSTM1* predicted expression is significantly associated with lower CHOL and LDL levels and is located 378 kb downstream from *SORT1* (Table 2). A study in north India found no significant difference in any lipid phenotype between individuals with *GSTM1* (−) and *GSTM1* (+), but did find that the *GSTM1* (−) genotype had a 2-fold increased risk of developing coronary artery disease in the cohort [60]. This gene has also been studied for association with atherosclerosis, and frequency of atherosclerosis in a *GSTM1* polymorphic group were found to be 1.2 times higher than those in the control group in a study from a Brazilian cohort [61].

In many studies, participants are asked to self-identify under a given label, such as the four race/ethnicity classifications used in MESA: “White, Caucasian”, “Chinese American”, “Black, African-American”, and “Hispanic”. These self-identifications are often not indicative of genetic ancestry, especially in admixed populations such as African-Americans and Hispanics due to a complex population history [32]. For example, even within self-identified regions in Latin America, populations can be heavily stratified (Fig 1 and S1 Fig). Our group recently created the first multi-ethnic transcriptomic prediction models for PrediXcan and observed that predictive performance was improved in cohorts of similar ancestry [14]. In our current analysis, we found *PSRC1* and *CETP* to be significantly associated with multiple phenotypes in MESA models, and these genes have been previously extensively studied in association with lipids [49]. The majority of HCHS/SoL results replicated in both GLGC and MESA even though there is a 50-fold difference in sample size (Table 2).

Recent studies have tested the portability of PrediXcan models across populations, notably between African and European populations. When predicting gene expression using GTEx models, the Yoruba population from West Africa had poorer accuracy compared to several European-ancestry populations tested [62]. Another study found similar results as both simulated and real African-American populations had poorer correlations of predicted versus observed gene expression than European populations [63]. Accuracy varied with both model population sample size and ancestry [63]. Both studies emphasized that a shared genetic architecture and ancestry between test and reference populations is imperative for accurate gene expression prediction and that further efforts in collection and creating multi-ethnic models are needed [62, 63].

In 2012, Stranger et al. observed significant sharing of eQTL effect sizes between Asians, European-admixed, and African subpopulations and suggested that the driving force behind the discovery of an eQTL in one population but not another is mainly due to allele frequency differences and not due to differences in absolute effect size [64]. Indeed, we recently showed differences in gene expression predictive performance are due to allele frequency differences between populations [14]. In MESA, we also showed that genetic correlation (rG) of eQTLs depends on shared ancestry proportions, ranging from rG=0.46 between AFA and CAU to rG=0.62 between HIS and CAU [14]. Thus, each population has shared and unique eQTL effects. Other than population-specific SNPs present in the MESA monocyte prediction models, SNPs that are both rare in European populations and common in Hispanic populations are not present in our analyses here and we continue to seek larger and more diverse gene expression data to improve inclusion of population-specific effects.

Another recent study showed that incorporating the use of local ancestry can improve eQTL mapping and gene expression prediction [65]. Future studies of lipid traits in larger Hispanic populations may include testing local ancestry portions individually in this population to see if less-studied ancestries, such as Native American and West African, are associated with SNPs and genes that have not been previously observed in European-majority cohorts. In our admixture mapping analysis to determine if significant GWAS signals originate from tracts of African or Native American ancestry within HCHS/SoL, we did not find any significant results, possibly due to the small non-admixed Native American reference cohort (n = 27) we used from the 1000 Genomes Project. This cohort is mainly Peruvian, which is likely not the best reference panel for the Native American component of most individuals in HCHS/SoL [66]. To have a higher power in these analyses, more samples, especially from Native American genomes and transcriptomes, are needed to integrate local ancestry into gene expression studies. By combining information from both local ancestry mapping and eQTL studies in non-European populations, we may better characterize and predict gene expression in admixed populations.

There is a dearth of diversity of genetic studies that greatly impacts the ability to apply the results of genetic studies to non-European populations. With many biobank-sized resources only including European data, this gap continues to grow [6]. Without data collection and proper models for non-European populations, there is less potential for accurate implementation of precision medicine. To fully characterize the impact of genetic variation between and within populations and allow all individuals to benefit from biomedical research, we must expand genetic studies to include individuals of diverse ancestries.

## Materials and Methods

This work was approved by the Loyola University Chicago Institutional Review Board (project number 2014).

### Phenotypic and genomic data

The analyses in HCHS/SoL and MESA were performed with whole genome genotypes from the database of Genotypes and Phenotypes (Table 4) [67]. The HCHS/SoL cohort is composed of self-identified Hispanic adults (18-74) recruited from four urban centers in the US: Chicago, IL; Miami, FL; the Bronx, NY; and San Diego, CA, using a previously described household-based sampling technique with written consent of the individual in their preferred language with data subsequently de-identified [7]. Within the collected lipid phenotypes, we rank-normalized CHOL, HDL, TRIG, and LDL to correct the skewing in the raw data. The majority of participants reported a self-identified region from Cuba (n = 1,987), Dominican Republic (n = 1,132), Mexico (n = 4,056), Puerto Rico (n = 1,984), Central America (n = 1,244), or South America (n = 707). Original genotypes were collected from a standard Illumina Omni2.5M array with 109,571 custom SNPs.

**Table 4.** Genotyped cohorts.

|  | HCHS | MESA |
| --- | --- | --- |
| Accession number | phs000810.v1.p1 | phs000209.v13.p3 |
| Pre-QC SNPs | 2,536,661 | 909,622 |
| Pre-QC individuals | 12,434 | 8,224 |
| Post-QC SNPs | 2,074,058 | 819,939 |
| Post-QC individuals | 12,235 | 8,224 |
| Post-imputation SNPs | 7,576,834 | 7,043,460 |
| Ind. with lipid pheno. | 11,104 | 3,855 |

Within the MESA cohort, individuals were recruited from six urban centers in the US: Baltimore, MD; Chicago, IL; Forsyth County, NC; Los Angeles County, CA; northern Manhattan, NY; and St. Paul, MN. Genotype data were collected from blood samples using an Affymetrix 6.0 SNP array and phenotype data from Exam 1 were used. MESA individuals include those that self-identified as “Black, African-American” (AFA, n = 613), “White, Caucasian” (CAU = 2,243), and “Hispanic” (HIS, n = 999). Cohorts underwent quality control and imputation as previously performed by our group [14].

Summary statistics from the Global Lipids Genetics Consortium were downloaded from http://csg.sph.umich.edu/willer/public/lipids2013/. This cohort is >95% European and contained 196,475 individuals. The original study had quantile-normalized phenotypes [5].

### Quality control

For quality control, we merged the two HCHS/SoL permission groups from dbGaP and ran all processes in PLINK. All of the genotypes are in genome build GRCh37/hg19. We removed individuals with genotyping rate < 99%. Additionally, we removed SNPs with failed heterozygosity (outside ± 3 standard deviations from the mean), with failed Hardy-Weinberg equilibrium (P < 1×10^−6^), and restricted analyses to SNPs on autosomes only. Unlike the typical quality control process for non-admixed populations, we did not remove principal component outliers nor did we remove individuals with >0.125 identity by descent in HCHS/SoL as this would have included >66% of the sample [68–70].

### Imputation, relationship inference, and principal component calculation

For imputation, we used the University of Michigan Imputation Server with EAGLE2 phasing [71]. The reference panel was 1000 Genomes Phase 3, using all populations due to the multi-continental nature of the cohort [72]. We then filtered imputation results to SNPs with *R*^2^ > 0.8 and minor allele frequency > 1% within PLINK. Quality control and imputation for MESA was performed by our group previously with the same parameters [14]. In MESA, each subpopulation was imputed separated and the post-imputation quality control procedures involved filtering by R^2^ > 0.8 in each subpopulation, combining the imputations, filtering by MAF > 0.01, and keeping all SNPs with genotyping rate > 0.99.

Within this cohort, there are potentially confounding amounts of genetic substructure present [73] (Fig 1). In structured and admixed cohorts, such as Hispanic and African-American populations, software such as KING are robust to complex population structure for relatedness and principal component calculations. [16]. We calculated pedigrees, then inferred relationships across all individuals in the cohort into a relationship matrix, and also calculated the first 20 principal components using KING, and we used the first 5 principal components in the rest of the analyses as performed previously within this cohort and since PCs > 5 explained less than 5% of the variance each (Fig 1 and S2 Fig) [19, 74].

### Genome-wide association study

We used the software GEMMA for the GWAS portion of the study. We ran separate univariate linear mixed models for each of the four phenotypes and included the relationship matrix as random effects. For the covariates used as fixed effects within the model, we included self-identified region, use of lipid or other cardiovascular medication, and the first 5 of 20 principal components as previously conducted in this cohort [16]. P values presented were calculated using the Wald test [10]. We restricted the analyses to SNPs in PrediXcan models as this was the main focus of the analyses, analyzing in total 1,770,809 SNPs for each phenotype. We performed multi-SNP-based conditional and joint association analysis using GWAS data in GCTA-COJO, and we included the genotype data to use the linkage disequilibrium calculations in the actual genotypes with a collinearity cutoff of 0.9 and a standard genome-wide significance threshold of 5×10^−8^ [20, 21]. We report these results and not base GWAS results to emphasize independent SNPs.

### Local ancestry inference

Since local ancestry inference for a population with three or more origins requires reference populations, we used Iberian from Spain (IBS) and Yoruba from Ibadan, Nigeria (YRI), both from 1000 Genomes Phase 3, as representatives for European and West African ancestry, respectively [72]. The European component of Hispanic populations has been previously identified as most similar to modern-day Iberian populations, and the African component has been identified as most similar to the Yoruba people [75].

In ADMIXTURE, we ran the 1000G American (AMR) populations with the Native American sequences, 1000G IBS, and 1000G YRI as reference populations, and kept 1000G AMR individuals with > 90% native American ancestry to use as a Native American reference panel for uniformity with 1000G (NAT), including 4 Mexicans from Los Angeles, USA and 23 Peruvians from Lima, Peru [76]. In total, our reference panel populations were IBS (n = 107), NAT (n = 27), and YRI (n = 108). We restricted analyses to chromosomes 16 and 19 due to their known importance in lipid traits.

To prepare our genotypes from local ancestry estimation, we used HAPI-UR to infer haplotypes with a maximum HMM window size of 64 [77]. We calculated local ancestry inference using RFMix in PopPhased mode, and due to our unequal reference population sizes, following the recommendation to use a minimum node size of 5 and a window size of 0.025, with all other options run as default [29].

### Admixture mapping

We converted the RFMix output to the three continental ancestry allelic estimations (NAT, IBS, and YRI). We tested each ancestry allele count for association with each phenotype to observe how local ancestry may be associated with each lipid trait using a univariate linear mixed model including relatedness as random effects and the first 5 principal components, lipid medications, and region as fixed effects [10]. Genome-wide admixture mapping significance was previously determined in HCHS/SoL at 5.7×10^−5^ [27].

### Imputed transcriptome-based association study

We used the software PrediXcan to produce predicted expression from genotype for both HCHS/SoL and MESA. The models we used include 44 tissue models from the GTEx V6 tissues with at least 70 individuals (85% European, 15% African-American) each, with an average of 5,179 genes predicted in each tissue [11, 13, 30]. We did not use GTEx V7 as the models only include individuals of European ancestry. We filtered the GTEx V6 results by removing red- and yellow-flagged false positive gene-tissue associations and our results only include gene-model combinations with green flags as described at http://predictdb.org/. The other set of PrediXcan models we used were made from monocyte gene expression in MESA [14]. Each of these five models had a training set of at least 233 individuals, including models of African-American (AFA), European (CAU), Hispanic (HIS), AFA and HIS (AFHI), and all populations combined (ALL). These models were retrieved from http://predictdb.org/, and all models in the analyses were filtered by protein coding genes, R^2^ > 0.01 and predictive performance P < 0.05. Individuals used to train the AFHI, ALL, and HIS models are also included in the MESA lipid trait replication cohort [14].

After the 49 predicted expression files were made using PrediXcan, we ran them as a pseudo-genotype in a univariate linear mixed model in GEMMA using the command -`notsnp`. This was performed in GEMMA rather than base PrediXcan as base PrediXcan does not have an option to include a relationship matrix, which is important in a heavily related and structured cohort like HCHS/SoL [10]. We again included the KING relationship matrix as random effects and the first five principal components, use of lipid medication, and self-identified region as fixed effects. P values presented were calculated using the Wald test.

Since we did not have the GLGC genotypes, we used the software S-PrediXcan, which is an extension of PrediXcan that takes GWAS summary statistics as input [13]. We ran S-PrediXcan on GLGC with all 44 GTEx tissue models and all 5 MESA monocyte models in different ethnicities. We considered significance in the discovery population, HCHS/SoL, as P < 3.1×10^−8^, 0.05/(all gene-model associations), and in the replication populations, MESA and GLGC, as P < 0.05 within the model. This significance threshold is conservative, since many tissues share eQTLs [31].

### Colocalization analysis

We performed a colocalization analysis by applying the software COLOC to the lipid GWAS results and eQTL data from GTEx and MESA to determine whether eQTLs within gene prediction models and GWAS hits were shared [15]. We subset the COLOC input to only contain SNPs within the predictor models of genes from the PrediXcan analyses due to computational restraints. A higher P4 probability (P > 0.5) indicates likely colocalized signals between an eQTL and a GWAS hit, especially in well-predicted genes with a high *R*^2^ value, while a high P3 probability indicates independent signals between an eQTL and a GWAS hit and a high P0, P1, or P2 indicates an unknown association [13]. Analyses were run using scripts from the S-PrediXcan manuscript at https://github.com/hakyimlab/summary-gwas-imputation/wiki/Running-Coloc [13].

## Acknowledgements

We would like to thank Peter Fiorica, Ryan Schubert, and Paul Okoro for their aid in reviewing the manuscript.

## Supporting information

**S1 Table. HCHS/SoL GCTA-COJO results for independent GWAS loci across all lipid phenotypes.**

**S2 Table. Top 1000 SNPs with most significant P values for admixture mapping.**

**S1 Fig. Genotypic principal component analysis in HCHS/SoL by self-identified regions.** PC1 vs. PC2 is plotted for each individual separated by their self-identified region. From previous observations and studies, Hispanic populations have multiple continental ancestries due to a previous history of colonization and slavery: African (bottom right, YRI), Native American (left, NAT), and European (top, EUR). Caribbean populations, such as the Cuban, Dominican, and Puerto Rican groups, tend to be mainly admixed between African and European, while mainland populations such as Mexican, Central American, and South American, tend to be mainly admixed between Native American and European.

**S2 Fig. Scree plot of the proportion of variance explained by the first 20 principal components.** Principal components 1, 2, and 3, explain 16.934%, 11.060%, and 5.494% of the variance, respectively. All other principal components explain < 5% of the variance each. All analyses used 5 PCs as fixed effects, as previously used in analyses of HCHS/SoL.

**S3 Fig. Quantile-quantile plots of the four lipid traits for GWAS results in HCHS/SoL.** Genomic control lambda values (*λ*) indicate little genome-wide deviation from the significance expectation line in any of the GWAS results.

**S4 Fig. Quantile-quantile plots of the four lipid traits for PrediXcan results in HCHS/SoL.** PrediXcan results for all 44 GTEx tissue models and 5 MESA monocyte populations are combined. Each point is a gene-tissue or gene-population association. Genomic control lambda values (*λ*) indicate little genome-wide deviation from the significance expectation line in any of the PrediXcan results.

## References

1. Mozaffarian D, Benjamin EJ, Go AS, Arnett DK, Blaha MJ, Cushman M, et al. Heart Disease and Stroke Statistics-2016 Update: A Report From the American Heart Association. Circulation. 2016;133(4):e38–360. doi:10.1161/CIR.0000000000000350.

2. Wu Y, Waite LL, Jackson AU, Sheu WHHH, Buyske S, Absher D, et al. Trans-Ethnic Fine-Mapping of Lipid Loci Identifies Population-Specific Signals and Allelic Heterogeneity That Increases the Trait Variance Explained. PLoS Genetics. 2013;9(3). doi:10.1371/journal.pgen.1003379.

3. Rodriguez CJ, Cai J, Swett K, González HM, Talavera GA, Wruck LM, et al. High Cholesterol Awareness, Treatment, and Control Among Hispanic/Latinos: Results From the Hispanic Community Health Study/Study of Latinos. Journal of the American Heart Association. 2015;4(7):1–10. doi:10.1161/JAHA.115.001867.

4. Popejoy AB, Fullerton SM. Genomics is failing on diversity. Nature. 2016;538(7624):161–164. doi:10.1038/538161a.

5. Willer CJ, Schmidt EM, Sengupta S, Peloso GM, Gustafsson S, Kanoni S, et al. Discovery and refinement of loci associated with lipid levels. Nature genetics. 2013;45(11):1274–83. doi:10.1038/ng.2797.

6. Martin AR, Kanai M, Kamatani Y, Okada Y, Neale BM, Daly MJ. Clinical use of current polygenic risk scores may exacerbate health disparities. Nature Genetics. 2019;51(4):584–591. doi:10.1038/s41588-019-0379-x.

7. LaVange LM, Kalsbeek WD, Sorlie PD, Avilés-Santa LM, Kaplan RC, Barnhart J, et al. Sample Design and Cohort Selection in the Hispanic Community Health Study/Study of Latinos. Annals of Epidemiology. 2010;20(8):642–649. doi:10.1016/j.annepidem.2010.05.006.

8. Andaleon A, Mogil LS, Wheeler HE. Gene-based association study for lipid traits in diverse cohorts implicates BACE1 and SIDT2 regulation in triglyceride levels. PeerJ. 2018;6(1):e4314. doi:10.7717/peerj.4314.

9. Graff M, Emery LS, Justice AE, Parra E, Below JE, Palmer ND, et al. Genetic architecture of lipid traits in the Hispanic community health study/study of Latinos. Lipids in Health and Disease. 2017;16(1):1–12. doi:10.1186/s12944-017-0591-6.

10. Zhou X, Stephens M. Genome-wide efficient mixed-model analysis for association studies. Nature Genetics. 2012;44(7):821–824. doi:10.1038/ng.2310.

11. Gamazon ER, Wheeler HE, Shah KP, Mozaffari SV, Aquino-Michaels K, Carroll RJ, et al. A gene-based association method for mapping traits using reference transcriptome data. Nature Genetics. 2015;47(9):1091–1098. doi:10.1038/ng.3367.

12. Wheeler HE, Shah KP, Brenner J, Garcia T, Aquino-Michaels K, Consortium G, et al. Survey of the Heritability and Sparse Architecture of Gene Expression Traits across Human Tissues. PLoS Genetics. 2016;12(11):e1006423. doi:10.1371/journal.pgen.1006423.

13. Barbeira AN, Dickinson SP, Bonazzola R, Zheng J, Wheeler HE, Torres JM, et al. Exploring the phenotypic consequences of tissue specific gene expression variation inferred from GWAS summary statistics. Nature Communications. 2018;9(1). doi:10.1038/s41467-018-03621-1.

14. Mogil LS, Andaleon A, Badalamenti A, Dickinson SP, Guo X, Rotter JI, et al. Genetic architecture of gene expression traits across diverse populations. PLoS Genetics. 2018;14(8):e1007586. doi:10.1371/journal.pgen.1007586.

15. Giambartolomei C, Vukcevic D, Schadt EE, Franke L, Hingorani AD, Wallace C, et al. Bayesian Test for Colocalisation between Pairs of Genetic Association Studies Using Summary Statistics. PLoS Genetics. 2014;10(5):e1004383. doi:10.1371/journal.pgen.1004383.

16. Conomos MP, Laurie CACCA, Stilp AM, Gogarten SM, McHugh CP, Nelson SC, et al. Genetic Diversity and Association Studies in US Hispanic/Latino Populations: Applications in the Hispanic Community Health Study/Study of Latinos. American Journal of Human Genetics. 2016;98(1):165–184. doi:10.1016/j.ajhg.2015.12.001.

17. Kramer HJ, Stilp AM, Laurie CC, Reiner AP, Lash J, Daviglus ML, et al. African AncestryâSpecific Alleles and Kidney Disease Risk in Hispanics/Latinos. Journal of the American Society of Nephrology. 2017;28(3):915–922. doi:10.1681/ASN.2016030357.

18. Belbin GM, Nieves-Colón MA, Kenny EE, Moreno-Estrada A, Gignoux CR. Genetic diversity in populations across Latin America: implications for population and medical genetic studies. Current Opinion in Genetics and Development. 2018;53:98–104. doi:10.1016/j.gde.2018.07.006.

19. Manichaikul A, Mychaleckyj JC, Rich SS, Daly K, Sale M, Chen WM. Robust relationship inference in genome-wide association studies. Bioinformatics. 2010;26(22):2867–2873. doi:10.1093/bioinformatics/btq559.

20. Yang J, Lee SH, Goddard ME, Visscher PM. GCTA: A tool for genome-wide complex trait analysis. American Journal of Human Genetics. 2011;88(1):76–82. doi:10.1016/j.ajhg.2010.11.011.

21. Yang J, Ferreira T, Morris AP, Medland SE, Madden PAFF, Heath AC, et al. Conditional and joint multiple-SNP analysis of GWAS summary statistics identifies additional variants influencing complex traits. Nature Genetics. 2012;44(4):369–375. doi:10.1038/ng.2213.

22. Klarin D, Damrauer SM, Cho K, Sun YV, Teslovich TM, Honerlaw J, et al. Genetics of blood lipids among ~300,000 multi-ethnic participants of the Million Veteran Program. Nature Genetics. 2018;50(11):1514–1523. doi:10.1038/s41588-018-0222-9.

23. Ferri GM, Brooks-Wilson AR, Kricker A, Staines A, Wu X, Habermann TM, et al. A genome-wide association study of marginal zone lymphoma shows association to the HLA region. Nature Communications. 2015;6(1):1–7. doi:10.1038/ncomms6751.

24. Pruim RJ, Welch RP, Sanna S, Teslovich TM, Chines PS, Gliedt TP, et al. LocusZoom: Regional visualization of genome-wide association scan results. Bioinformatics. 2011;27(13):2336–2337. doi:10.1093/bioinformatics/btq419.

25. Sabatti C, Service SK, Hartikainen AL, Pouta A, Ripatti S, Brodsky J, et al. Genome-wide association analysis of metabolic traits in a birth cohort from a founder population. Nature Genetics. 2009;41(1):35–46. doi:10.1038/ng.271.

26. Zhou X. a Unified Framework for Variance Component Estimation With Summary Statistics in Genome-Wide Association Studies. The annals of applied statistics. 2017;11(4):2027–2051. doi:10.1214/17-AOAS1052.

27. Brown LA, Sofer T, Stilp AM, Baier LJ, Kramer HJ, Masindova I, et al. Admixture Mapping Identifies an Amerindian Ancestry Locus Associated with Albuminuria in Hispanics in the United States. Journal of the American Society of Nephrology. 2017;28(7):2211–2220. doi:10.1681/ASN.2016091010.

28. Sofer T, Baier LJ, Browning SR, Thornton TA, Talavera GA, Wassertheil-Smoller S, et al. Admixture mapping in the Hispanic Community Health Study/Study of Latinos reveals regions of genetic associations with blood pressure traits. PLoS ONE. 2017;12(11):1–15. doi:10.1371/journal.pone.0188400.

29. Maples BK, Gravel S, Kenny EE, Bustamante CD. RFMix: A discriminative modeling approach for rapid and robust local-ancestry inference. American Journal of Human Genetics. 2013;93(2):278–288. doi:10.1016/j.ajhg.2013.06.020.

30. Ardlie KG, DeLuca DS, Segrè AV, Sullivan TJ, Young TR, Gelfand ET, et al. The Genotype-Tissue Expression (GTEx) pilot analysis: Multitissue gene regulation in humans. Science. 2015;348(6235):648–660. doi:10.1126/science.1262110.

31. Barbeira AN, Pividori MD, Zheng J, Wheeler HE, Nicolae DL, Im HK. Integrating predicted transcriptome from multiple tissues improves association detection. PLoS Genetics. 2019;15(1):e1007889. doi:10.1371/journal.pgen.1007889.

32. Bild DE, Bluemke DA, Burke GL, Detrano R, Diez Roux AV, Folsom AR, et al. Multi-Ethnic Study of Atherosclerosis: Objectives and design. American Journal of Epidemiology. 2002;156(9):871–881. doi:10.1093/aje/kwf113.

33. Zhou L, He M, Mo Z, Wu C, Yang H, Yu D, et al. A genome wide association study identifies common variants associated with lipid levels in the Chinese population. PLoS ONE. 2013;8(12):e82420. doi:10.1371/journal.pone.0082420.

34. MacArthur J, Bowler E, Cerezo M, Gil L, Hall P, Hastings E, et al. The new NHGRI-EBI Catalog of published genome-wide association studies (GWAS Catalog). Nucleic Acids Research. 2017;45(D1):D896–D901. doi:10.1093/nar/gkw1133.

35. Thompson JF, Lira ME, Durham LK, Clark RW, Bamberger MJ, Milos PM. Polymorphisms in the CETP gene and association with CETP mass and HDL levels. Atherosclerosis. 2003;167(2):195–204. doi:10.1016/S0021-9150(03)00005-4.

36. Petty LE, Highland HM, Gamazon ER, Hu H, Karhade M, Chen HH, et al. Functionally oriented analysis of cardiometabolic traits in a trans-ethnic sample. Human Molecular Genetics. 2019;28(7):1212–1224. doi:10.1093/hmg/ddy435.

37. Feingold KR, Grunfeld C. Introduction to Lipids and Lipoproteins. South Dartmouth, MA: Endotext [Internet]; 2000. Available from: https://www.ncbi.nlm.nih.gov/books/NBK305896/.

38. Luna A, Nicodemus KK. snp.plotter: an R-based SNP/haplotype association and linkage disequilibrium plotting package. Bioinformatics. 2007;23(6):774–776. doi:10.1093/bioinformatics/btl657.

39. Liu B, Gloudemans MJ, Rao AS, Ingelsson E, Montgomery SB. Abundant associations with gene expression complicate GWAS follow-up. Nature Genetics. 2019;51(5):768–769. doi:10.1038/s41588-019-0404-0.

40. Strong A, Patel K, Rader DJ. Sortilin and lipoprotein metabolism. Current Opinion in Lipidology. 2014;25(5):350–357. doi:10.1097/mol.0000000000000110.

41. Guo K, Hu L, Xi D, Zhao J, Liu J, Luo T, et al. PSRC1 overexpression attenuates atherosclerosis progression in apoE−/− mice by modulating cholesterol transportation and inflammation. Journal of Molecular and Cellular Cardiology. 2018;116(January):69–80. doi:10.1016/j.yjmcc.2018.01.013.

42. Arvind P, Nair J, Jambunathan S, Kakkar VV, Shanker J. CELSR2-PSRC1-SORT1 gene expression and association with coronary artery disease and plasma lipid levels in an Asian Indian cohort. Journal of Cardiology. 2014;64(5):339–346. doi:10.1016/j.jjcc.2014.02.012.

43. Barter PJ, Brewer HB, Chapman MJ, Hennekens CH, Rader DJ, Tall AR. Cholesteryl ester transfer protein: A novel target for raising HDL and inhibiting atherosclerosis. Arteriosclerosis, Thrombosis, and Vascular Biology. 2003;23(2):160–167. doi:10.1161/01.ATV.0000054658.91146.64.

44. de Grooth GJ, Klerkx AHEM, Stroes ESG, Stalenhoef AFH, Kastelein JJP, Kuivenhoven JA. A review of CETP and its relation to atherosclerosis. Journal of Lipid Research. 2004;45(11):1967–1974. doi:10.1194/jlr.r400007-jlr200.

45. Kosmas CE, Dejesus E, Rosario D, Vittorio TJ. CETP inhibition: Past failures and future hopes. Clinical Medicine Insights: Cardiology. 2016;10:37–42. doi:10.4137/CMC.S32667.

46. Tall AR, Rader DJ. Trials and Tribulations of CETP Inhibitors. Circulation Research. 2018;122(1):106–112. doi:10.1161/CIRCRESAHA.117.311978.

47. Wainberg M, Sinnott-armstrong N, Mancuso N, Barbeira AN, Knowles DA, Golan D, et al. Opportunities and challenges for transcriptome-wide association studies. Nature Genetics. 2019;51(April). doi:10.1038/s41588-019-0385-z.

48. Musunuru K, Strong A, Frank-Kamenetsky M, Lee NE, Ahfeldt T, Sachs KV, et al. From noncoding variant to phenotype via SORT1 at the 1p13 cholesterol locus. Nature. 2010;466(7307):714. doi:10.1038/NATURE09266.

49. Below JE, Parra EJ, Gamazon ER, Torres J, Krithika S, Candille S, et al. Meta-analysis of lipid-traits in Hispanics identifies novel loci, population-specific effects, and tissue-specific enrichment of eQTLs. Scientific Reports. 2016;6:19429. doi:10.1038/srep19429.

50. Surakka I, Horikoshi M, Mägi R, Sarin AP, Mahajan A, Lagou V, et al. The impact of low-frequency and rare variants on lipid levels. Nature Genetics. 2015;47(6):589–597. doi:10.1038/ng.3300.

51. Paththinige CS, Sirisena ND, Dissanayake VHWW. Genetic determinants of inherited susceptibility to hypercholesterolemia - a comprehensive literature review. Lipids in Health and Disease. 2017;16(1):1–22. doi:10.1186/s12944-017-0488-4.

52. Weissglas-Volkov D, Aguilar-Salinas CA, Nikkola E, Deere KA, Cruz-Bautista I, Arellano-Campos O, et al. Genomic study in Mexicans identifies a new locus for triglycerides and refines European lipid loci. Journal of Medical Genetics. 2013;50(5):298–308. doi:10.1136/jmedgenet-2012-101461.

53. Beecham AH, Patsopoulos NA, Xifara DK, Davis MF, Kemppinen A, Cotsapas C, et al. Analysis of immune-related loci identifies 48 new susceptibility variants for multiple sclerosis. Nature Genetics. 2013;45(11):1353–1362. doi:10.1038/ng.2770.

54. Langefeld CD, Ainsworth HC, Graham DSC, Kelly JA, Comeau ME, Marion MC, et al. Transancestral mapping and genetic load in systemic lupus erythematosus. Nature Communications. 2017;8(May). doi:10.1038/ncomms16021.

55. Kimura S, Wang Ky, Yamada S, Guo X, Nabeshima A, Noguchi H, et al. CCL22/Macrophage-derived Chemokine Expression in Apolipoprotein E-deficient Mice and Effects of Histamine in the Setting of Atherosclerosis. Journal of Atherosclerosis and Thrombosis. 2015;22(6):599–609. doi:10.5551/jat.27417.

56. Kimura S, Tanimoto A, Wang KY, Shimajiri S, Guo X, Tasaki T, et al. Expression of macrophage-derived chemokine (CCL22) in atherosclerosis and regulation by histamine via the H2 receptor. Pathology International. 2012;62(10):675–683. doi:10.1111/j.1440-1827.2012.02854.x.

57. Rosenson RS, Brewer HB, Ansell BJ, Barter P, Chapman MJ, Heinecke JW, et al. Dysfunctional HDL and atherosclerotic cardiovascular disease. Nature Reviews Cardiology. 2016;13(1):48–60. doi:10.1038/nrcardio.2015.124.

58. Fotis L, Agrogiannis G, Vlachos IS, Pantopoulou A, Margoni A, Kostaki M, et al. Intercellular adhesion molecule (ICAM)-1 and vascular cell adhesion molecule (VCAM)-1 at the early stages of atherosclerosis in a rat model. In Vivo. 2018;26(2):243–250.

59. Tabet F, Vickers KC, Cuesta Torres LF, Wiese CB, Shoucri BM, Lambert G, et al. HDL-transferred microRNA-223 regulates ICAM-1 expression in endothelial cells. Nature Communications. 2014;5:1–14. doi:10.1038/ncomms4292.

60. Bhatti JS, Vijayvergiya R, Singh B, Bhatti GK. Genetic susceptibility of glutathione S-transferase genes (GSTM1/T1 and P1) to coronary artery disease in Asian Indians. Annals of Human Genetics. 2018;82(6):448–456. doi:10.1111/ahg.12274.

61. Rodrigues DA, Martins JVM, E Silva KSF, Costa IR, Lagares MH, Campedelli FL, et al. GSTM1 polymorphism in patients with clinical manifestations of atherosclerosis. Genetics and Molecular Research. 2017;16(1):1–9. doi:10.4238/gmr16019101.

62. Mikhaylova AV, Thornton TA. Accuracy of Gene Expression Prediction From Genotype Data With PrediXcan Varies Across and Within Continental Populations. Frontiers in Genetics. 2019;10(April):1–10. doi:10.3389/fgene.2019.00261.

63. Keys K, Mak ACY, White MJ, Eckalbar WL, Dahl AW, Mefford J, et al. On the cross-population portability of gene expression prediction models. bioRxiv. 2019;doi:https://doi.org/10.1101/552042.

64. Stranger BE, Montgomery SB, Dimas AS, Parts L, Stegle O, Ingle CE, et al. Patterns of Cis regulatory variation in diverse human populations. PLoS Genetics. 2012;8(4):e1002639. doi:10.1371/journal.pgen.1002639.

65. Zhong Y, Perera MA, Gamazon ER. On Using Local Ancestry to Characterize the Genetic Architecture of Human Traits: Genetic Regulation of Gene Expression in Multiethnic or Admixed Populations. The American Journal of Human Genetics. 2019;104(6):1097–1115. doi:10.1016/J.AJHG.2019.04.009.

66. Browning SR, Grinde K, Plantinga A, Gogarten SM, Stilp AM, Kaplan RC, et al. Local Ancestry Inference in a Large US-Based Hispanic/Latino Study: Hispanic Community Health Study/Study of Latinos (HCHS/SOL). G3: Genes, Genomes, Genetics. 2016;6(6):1525–1534. doi:10.1534/g3.116.028779.

67. Mailman MD, Feolo M, Jin Y, Kimura M, Tryka K, Bagoutdinov R, et al. The NCBI dbGaP database of genotypes and phenotypes. Nature Genetics. 2007;39(10):1181–1186. doi:10.1038/ng1007-1181.

68. Purcell S, Neale B, Todd-Brown K, Thomas L, Ferreira MARR, Bender D, et al. PLINK: A Tool Set for Whole-Genome Association and Population-Based Linkage Analyses. The American Journal of Human Genetics. 2007;81(3):559–575. doi:10.1086/519795.

69. Turner S, Armstrong LL, Bradford Y, Carlson CS, Dana C, Crenshaw AT, et al. Quality control procedures for genome wide association studies. Current Proceedings in Human Genetics. 2011;68(1):1–24. doi:10.1002/0471142905.hg0119s68.Quality.

70. Chang CC, Chow CC, Tellier LC, Vattikuti S, Purcell SM, Lee JJ. Second-generation PLINK: rising to the challenge of larger and richer datasets. GigaScience. 2015;4(7):1–16. doi:10.1186/s13742-015-0047-8.

71. Das S, Forer L, Schönherr S, Sidore C, Locke AE, Kwong A, et al. Next-generation genotype imputation service and methods. Nature Genetics. 2016;48(10):1284–1287. doi:10.1038/ng.3656.

72. Auton A, Abecasis GR, Altshuler DM, Durbin RM, Abecasis GR, Bentley DR, et al. A global reference for human genetic variation. Nature. 2015;526(7571):68–74. doi:10.1038/nature15393.

73. Daviglus ML, Talavera GA, Avilés-Santa ML, Allison M, Cai J, Criqui MH, et al. Prevalence of major cardiovascular risk factors and cardiovascular diseases among Hispanic/Latino individuals of diverse backgrounds in the United States. JAMA. 2012;308(17):1775–1784. doi:10.1001/jama.2012.14517.

74. Conomos MP, Miller MB, Thornton TA. Robust inference of population structure for ancestry prediction and correction of stratification in the presence of relatedness. Genetic Epidemiology. 2015;39(4):276–293. doi:10.1002/gepi.21896.

75. Moreno-Estrada A, Gravel S, Zakharia F, McCauley JL, Byrnes JK, Gignoux CR, et al. Reconstructing the Population Genetic History of the Caribbean. PLoS Genetics. 2013;9(11). doi:10.1371/journal.pgen.1003925.

76. Alexander DH, Novembre J, Lange K. Fast model-based estimation of ancestry in unrelated individuals. Genome Research. 2009;19(9):1655–1664. doi:10.1101/gr.094052.109.

77. Williams ALL, Patterson N, Glessner J, Hakonarson H, Reich D. Phasing of many thousands of genotyped samples. American Journal of Human Genetics. 2012;91(2):238–251. doi:10.1016/j.ajhg.2012.06.013.

