## Supplementary figures and images for "Genetically regulated gene expression underlies lipid traits in Hispanic cohorts"

### S1 Fig.

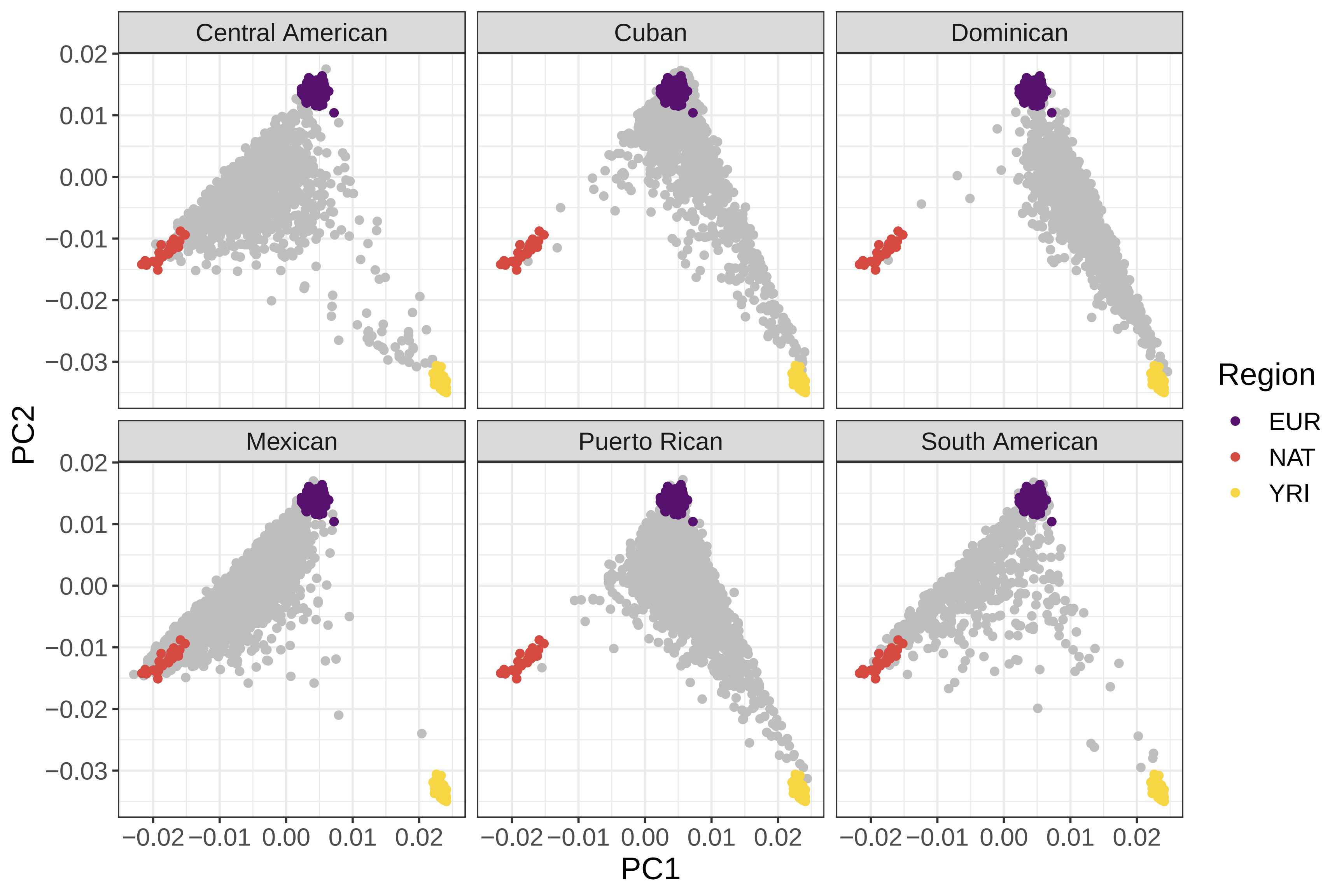

### S2 Fig.

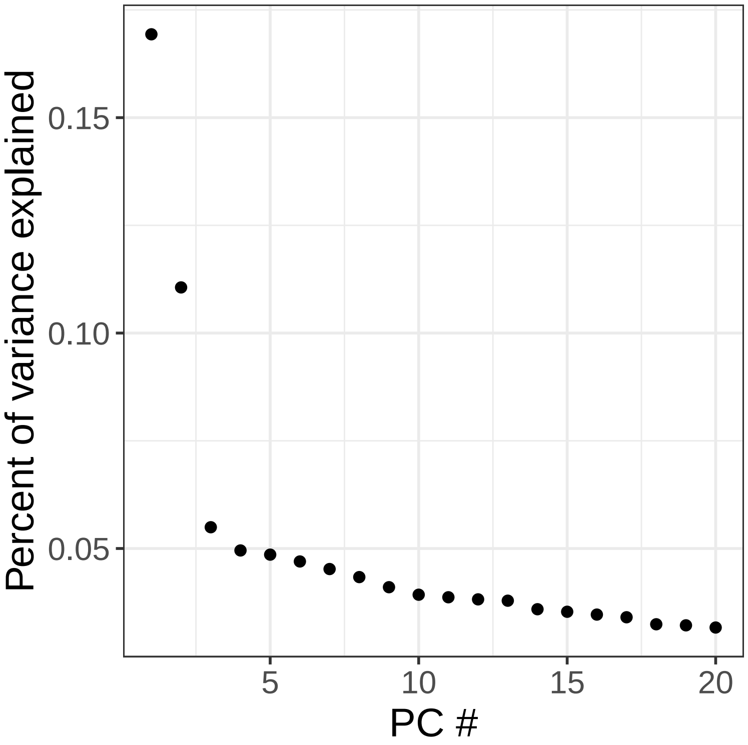

### S3 Fig.

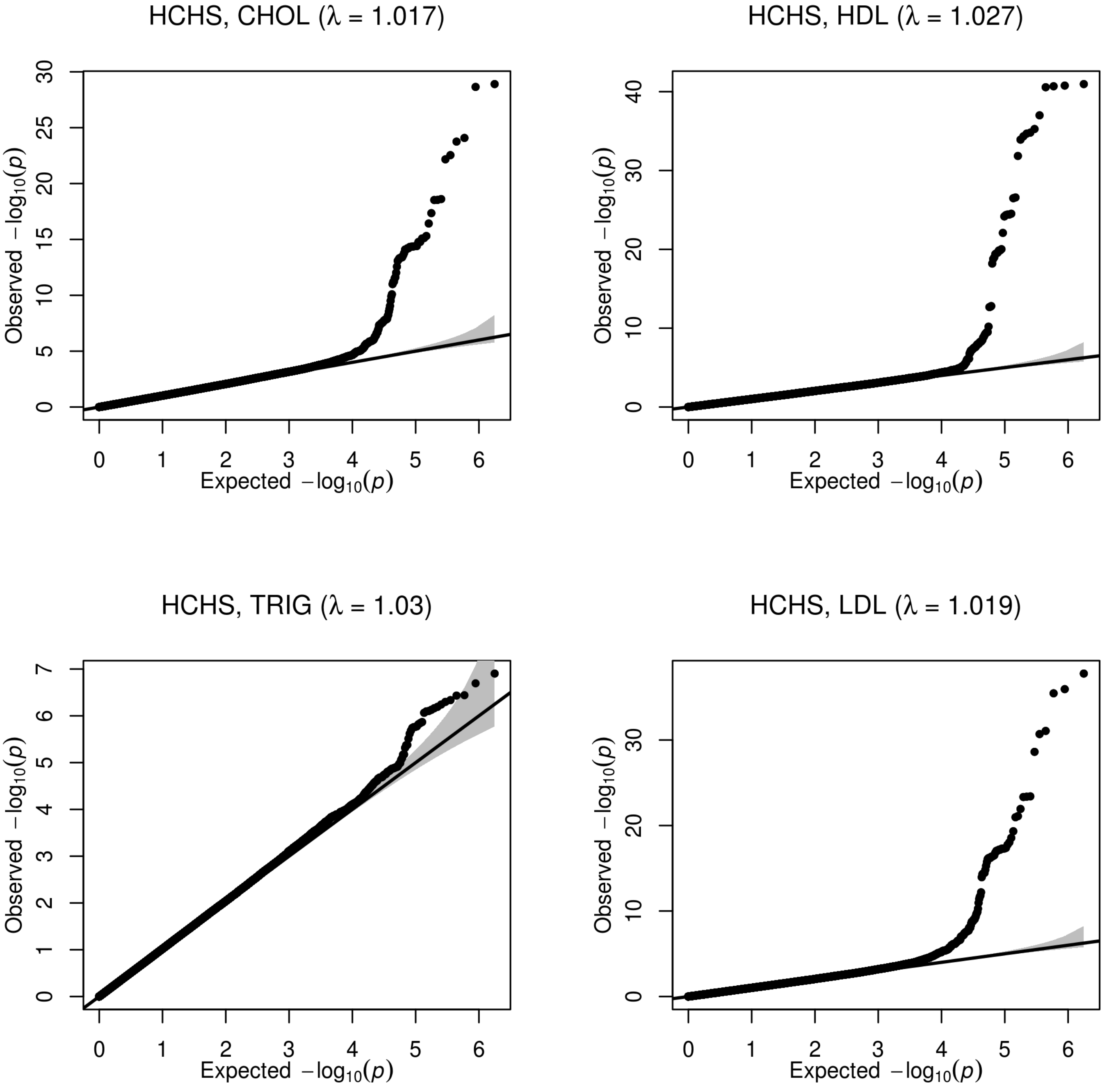

### S4 Fig.

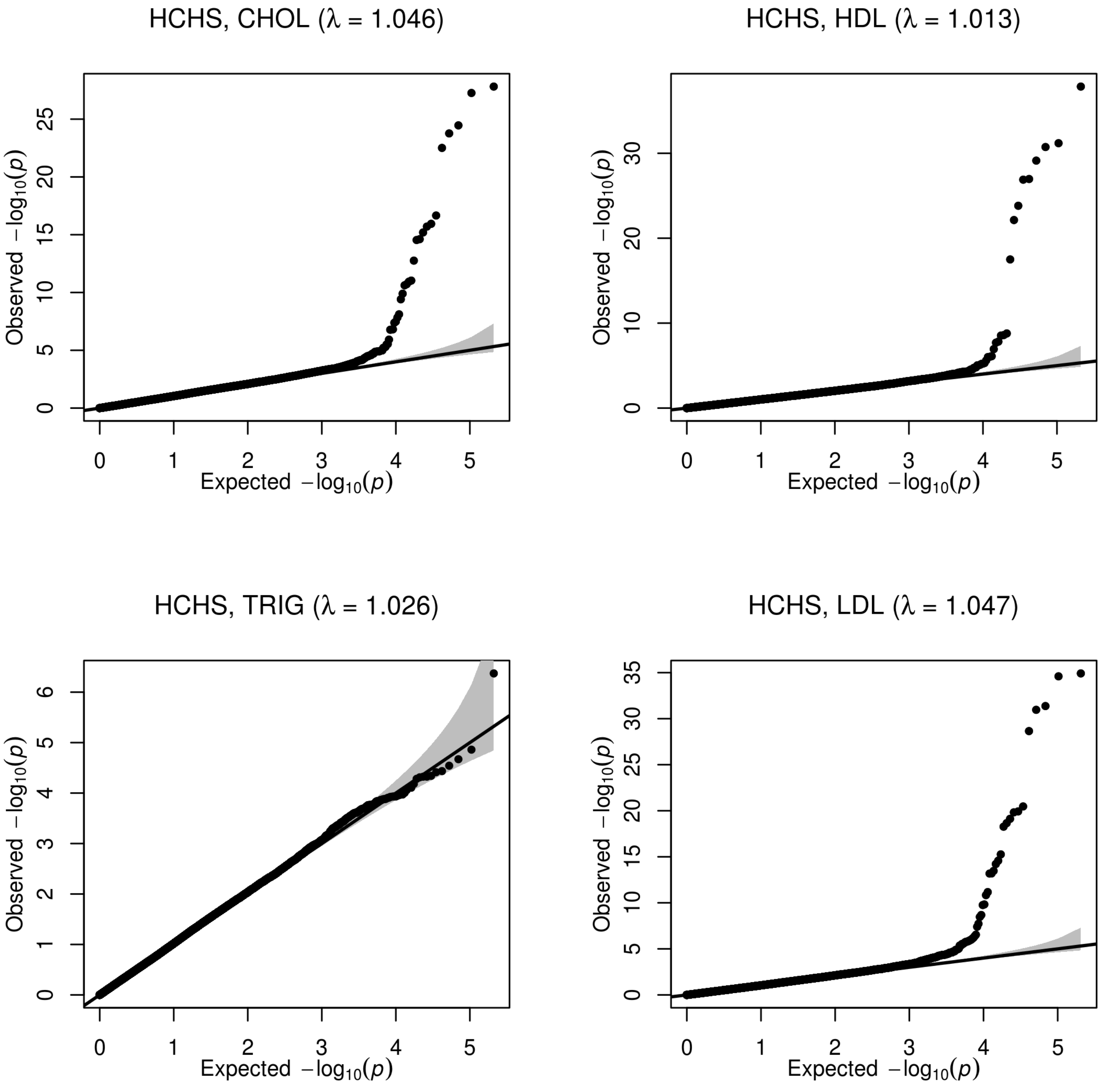
